# A Pre-Bilaterian Origin of Phototransduction Genes and Photoreceptor Cells

**DOI:** 10.1101/2025.02.09.637293

**Authors:** Alessandra Aleotti, Flaviano Giorgini, Roberto Feuda

## Abstract

The evolution of vision is a major novelty in animals, playing a fundamental role in developing complex behaviours. Vision initiates with a light-triggered phototransduction cascade occurring in photoreceptor cells (PRCs). The two main PRC types, ciliary and rhabdomeric, employ both specific and common genes for phototransduction. Despite being crucial for vision, the origin and evolution of photoreceptor cells and phototransduction pathways remain unclear.

Using phylogenetic methods, we studied the evolution of all phototransduction genes, elucidating their gene duplication patterns in over 80 species, including non-bilaterian metazoans and other eukaryotes. Then, we investigated the expression of phototransduction genes in available single-cell RNA-sequencing data from various animals, including non-bilaterians. By using phototransduction genes as markers, we identified putative photoreceptor-like cells across animals and compared their regulatory toolkits.

Gene families encoding phototransduction components are generally ancient, predating the origin of vision. However, many phototransduction genes originated in the metazoan stem group. Moreover, putative photoreceptor cells identified in non-bilaterians appeared to express some but not all components of the two well-characterised phototransduction pathways, suggesting potential lineage-specific components involved in phototransduction. Finally, we identified conserved expression of certain transcription factors in putative PRCs in non-bilaterians, suggesting the homology of PRCs.

## Introduction

The ability to detect light is fundamental for modulating interactions between animals and their environment (Nilsson 2009; Nilsson 2013). At the molecular level, this process begins with a photon of light activating an opsin (Terakita 2005), which triggers a chain of molecular reactions within the cell, culminating in electrical signalling. Two alternative phototransduction cascades specific to the different photoreceptor cells—ciliary and rhabdomeric—have been described in animals (Yau and Hardie 2009). In the rhabdomeric phototransduction cascade, extensively investigated in the fruit fly *Drosophila melanogaster* (**Figure S1A and Table S1**), opsin activates a Gq-type G protein. The alpha subunit of this G protein detaches and activates phospholipase C beta, initiating a phosphoinositide cascade. This results in the opening of transient receptor potential (TRP) and TRP-like (TRPL) channels, leading to cell depolarization (Wang and Montell 2007; Hardie and Juusola 2015). In the ciliary phototransduction cascade, extensively investigated in vertebrates (**Figure S1B and Table S1**), opsin activates transducin (Gt), a Gi/o-type G protein that activates phosphodiesterase 6 (PDE6), which hydrolyses cyclic GMP. The drop in cGMP levels causes cyclic nucleotide-gated ion channels (CNGCs) to close, resulting in cell hyperpolarization (Lamb 2020).

Although ciliary photoreceptors are predominantly found in deuterostomes, and rhabdomeric photoreceptors in protostomes, both types have been described in both bilaterian animals (Horridge 1964; Hattar et al. 2002; Arendt 2003; Nordström et al. 2003; Arendt et al. 2004; Kozmik et al. 2008; Passamaneck et al. 2011; Ullrich-Lüter et al. 2011; Jékely et al. 2015; Tamm 2016; von Döhren and Bartolomaeus 2018; Picciani et al. 2018; Valencia et al. 2021; Brodrick and Jékely 2023). While strong evidence suggests that these phototransduction cascades and their associated PRCs were present in the last common bilaterian ancestor and studies have described PRCs and associated gene in cnidarians (Nordström et al. 2003; Kozmik et al. 2008; Plachetzki et al. 2010; Mason et al. 2012), their presence in ctenophores, placozoans, and sponges remains uncertain. This raises two questions: When did the phototransduction cascade genes evolve, and when did they begin to operate in the same cell? Additionally, can we identify PRCs in non-bilaterian animals, and how are they related to those in bilaterians?

To address these questions, we investigated the distribution of each phototransduction component gene family (28 in total) in 86 eukaryotic species, focusing in early-branching animals and their close relatives, using state-of-the-art phylogenetic methods to infer duplication patterns. Next, by analysing existing single-cell RNA-sequencing data and using the co-expression of phototransduction genes, we identified putative PRC cells across various animal species and uncovered the co-expressed transcription factors (TFs). Our results suggest that most genes required for phototransduction evolved in the metazoan stem group from expansions within ancient gene families. Furthermore, we characterised the regulatory programs in PRC-like cells in non-bilaterians, uncovering some degree of conservation across all animals. Overall, these findings support a model in which the regulatory program of PRCs evolved by integrating a pre-existing core photoreceptor machinery under the regulation of the same TFs.

## Results

### Most phototransduction genes derive from metazoan-specific expansions in eukaryotic gene families

Initially, we analysed the phylogenetic distribution and evolutionary history of ciliary and rhabdomeric phototransduction proteins. We mined a dataset comprising 86 species (**Table S2**), including 25 non-bilaterian species and close relatives of animals (8 choanoflagellates and 5 other holozoans), and identified homologues of phototransduction genes using a combination of sequence similarity and protein motif analyses (see Materials and Methods). Importantly, several components required for the ciliary and rhabdomeric phototransduction cascades belong to large protein families that are involved in many biological processes beyond phototransduction. For example, G alpha proteins are required in numerous cell-signalling pathways, consequently, there are several paralog groups belonging to different subfamilies (Krishnan et al. 2015; Lokits et al. 2018). However, rhabdomeric phototransduction requires specifically the Gq type proteins, while ciliary phototransduction in humans utilises a Gi/o-type proteins (Yau and Hardie 2009). Similarly, the calmodulin family is involved in calcium-sensing in many biological processes throughout eukaryotes and is a vast family including both calmodulin and calmodulin-like members (Mohanta et al. 2017; Villalobo et al. 2019), with only one paralogue involved in phototransduction (Hardie 2012; Barret et al. 2023). To discriminate between orthologous genes and other paralogues within each gene family, we constructed maximum likelihood phylogenetic trees and performed gene tree-to-species tree reconciliations using GeneRax (Morel et al. 2020) (see Methods for details).

Our results indicate that in almost all cases, identifying one-to-one orthologues across all metazoans was impossible, as many genes underwent lineage-specific duplications (Figures S2-S29; all raw output files can be found on the FigShare repository: https://doi.org/10.25392/leicester.data.c.7284370). Furthermore, to account for uncertainty in early metazoan phylogeny (Ryan et al. 2013; Moroz et al. 2014; Pisani et al. 2015; Whelan et al. 2015; Feuda et al. 2017; Simion et al. 2017; Whelan et al. 2017; Kapli and Telford 2020; Giacomelli et al. 2022; Schultz et al. 2023), the gene tree-to-species tree reconciliations were performed with both ctenophore-first and sponge-first scenarios. A comparison of the total number of events (duplications and losses) revealed that, overall, there were no major differences between the two scenarios (**Table S3**).

Six gene families are common to both phototransduction pathways (Table S1). The distribution of homologous genes suggests that G beta, calmodulin, and GRK are present in all eukaryotes, arrestin in Holozoa, while opsin and G gamma are animal-specific. However, the gene tree-to-species tree reconciliation indicates that within the broad families, the subgroups involved in the phototransduction cascade are present either strictly in animals, as in the case of opsins, G gamma, and arrestin, or within Holozoa (G beta, calmodulin, GRK) (Figure 1A and Supplementary Figures S2-S7).

**Figure 1.**
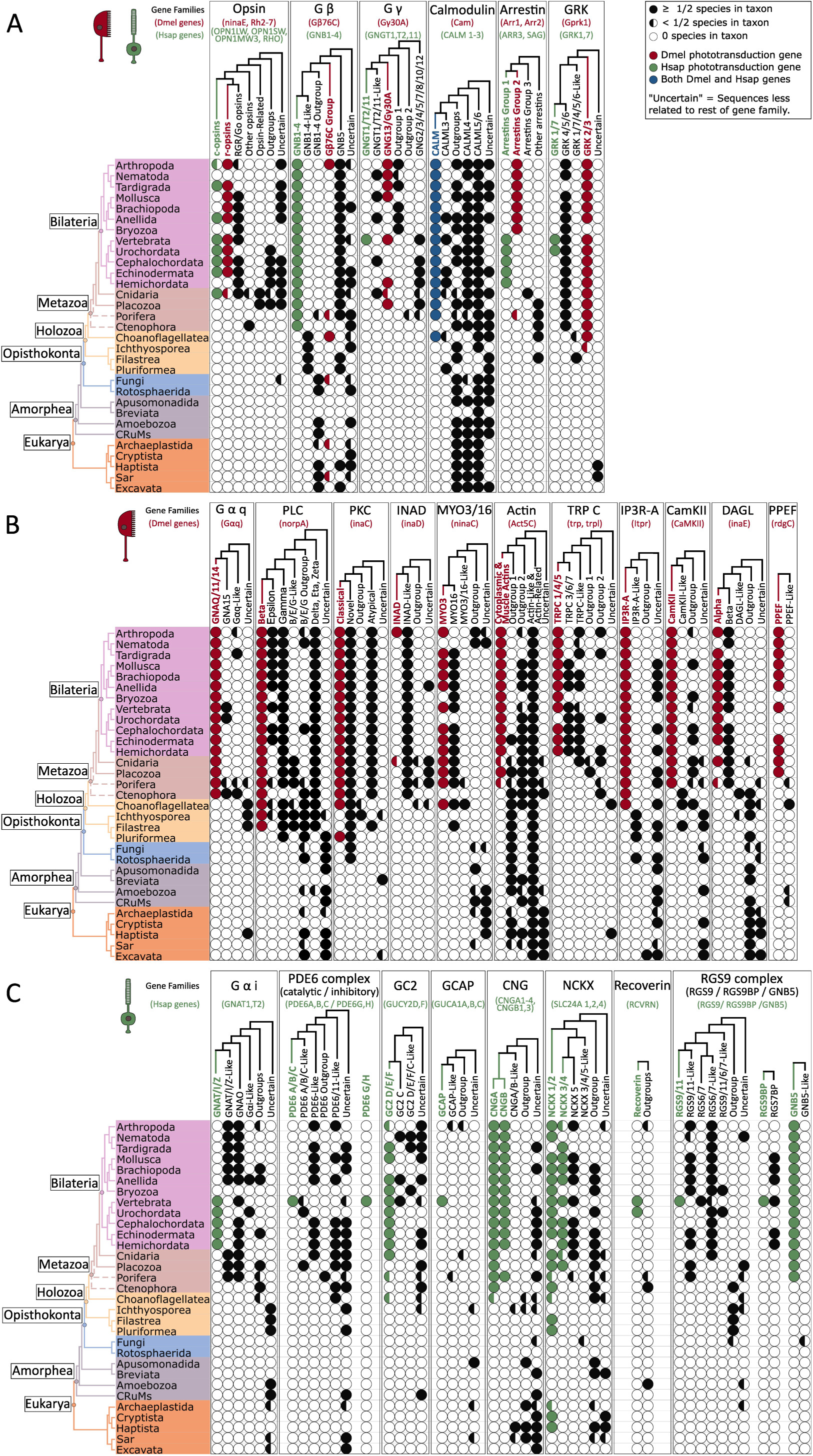
Evolutionary history of phototransduction components’ gene families and distribution in eukaryotes. The evolution of each gene family for all common (**A**), rhabdomeric-specific (**B**) and ciliary-specific (**C**) components is reconstructed (gene trees at the top of each section), and their distribution mapped across major groups of Eukarya (species tree on the left of each section). Presence is indicated with a fully-coloured circle if at least half of the species examined within the clade possess the gene or with a half-coloured circle if less than half of them possess it. Within each gene family the subfamilies of interest containing the *D. melanogaster* (red), *H. sapiens* (green) or both (blue) gene(s) known to function in the phototransduction pathway(s) are highlighted. In a few gene families, some sequences with poor annotation appear very distantly related to the group of interest according to the phylogeny. These clades are labelled as “Uncertain”. In fact, they could represent genuinely distantly related members of the gene family since they were retrieved during data mining and retained during the pipeline. However, it cannot be excluded that they might rather belong to a different gene family.

Homologues of rhabdomeric phototransduction components are distributed throughout eukaryotes, except for INAD, which appears to be restricted to animals and choanoflagellates. The subgroups involved in phototransduction are instead animal-specific in seven out of eleven cases, with the remaining four families being holozoan-specific (Figure 1B and Supplementary Figures S8-S18).

Most of the ciliary-specific extended gene families are also widely distributed with homologues present throughout eukaryotes for nine families and only two families being animal-specific (**Figure 1C and Supplementary Figures S19-S29**). The subgroups involved in phototransduction are conversely mostly animal-specific (eight out of eleven). Two components are also present in Holozoa, while only NCKX, a sodium-calcium-potassium exchanger involved in numerous other pathways (Altimimi and Schnetkamp 2007), is present throughout eukaryotes. Interestingly, our results suggest that ciliary phototransduction genes exhibit a patchy distribution within animals, unlike common and rhabdomeric components, which are distributed across all or most animal phyla. Vertebrates are the only group containing all the genes for ciliary components. This indicates that some components of the ciliary pathway are likely vertebrate innovations, while other components are more ancient and represent the core part of the cascade.

Overall, our results indicate that while homologues of most phototransduction genes are widely distributed in eukaryotes, recent duplication events expanded the gene families, giving rise to numerous subgroups, including key genes for phototransduction. These waves of duplications typically occurred within holozoans, especially at the stem of Metazoa. Furthermore, while rhabdomeric phototransduction genes are generally evenly distributed throughout Metazoa, ciliary genes are frequently vertebrate-specific, indicating more recent specialisations.

### The Evolution of Photoreceptor Cell Identity in Metazoans

Next, we used ciliary and rhabdomeric phototransduction cascade genes to identify putative photoreceptor cells (PRCs) in animals. To achieve this, we capitalised on our phylogenetic results and combined them with available single-cell RNA-sequencing data. We focused on twelve species spanning the metazoans, particularly non-bilaterian animals (**Figure S30**). To overcome technical differences in single-cell RNA-seq data, such as the dropout rate, we used the MetaCell pipeline (Baran et al. 2019) to identify “metacells” or cell states. We further identified those with a PRC-like profile (**Figure 2**) using a ranking system based on the expression of phototransduction genes (see methods).

**Figure 2.**
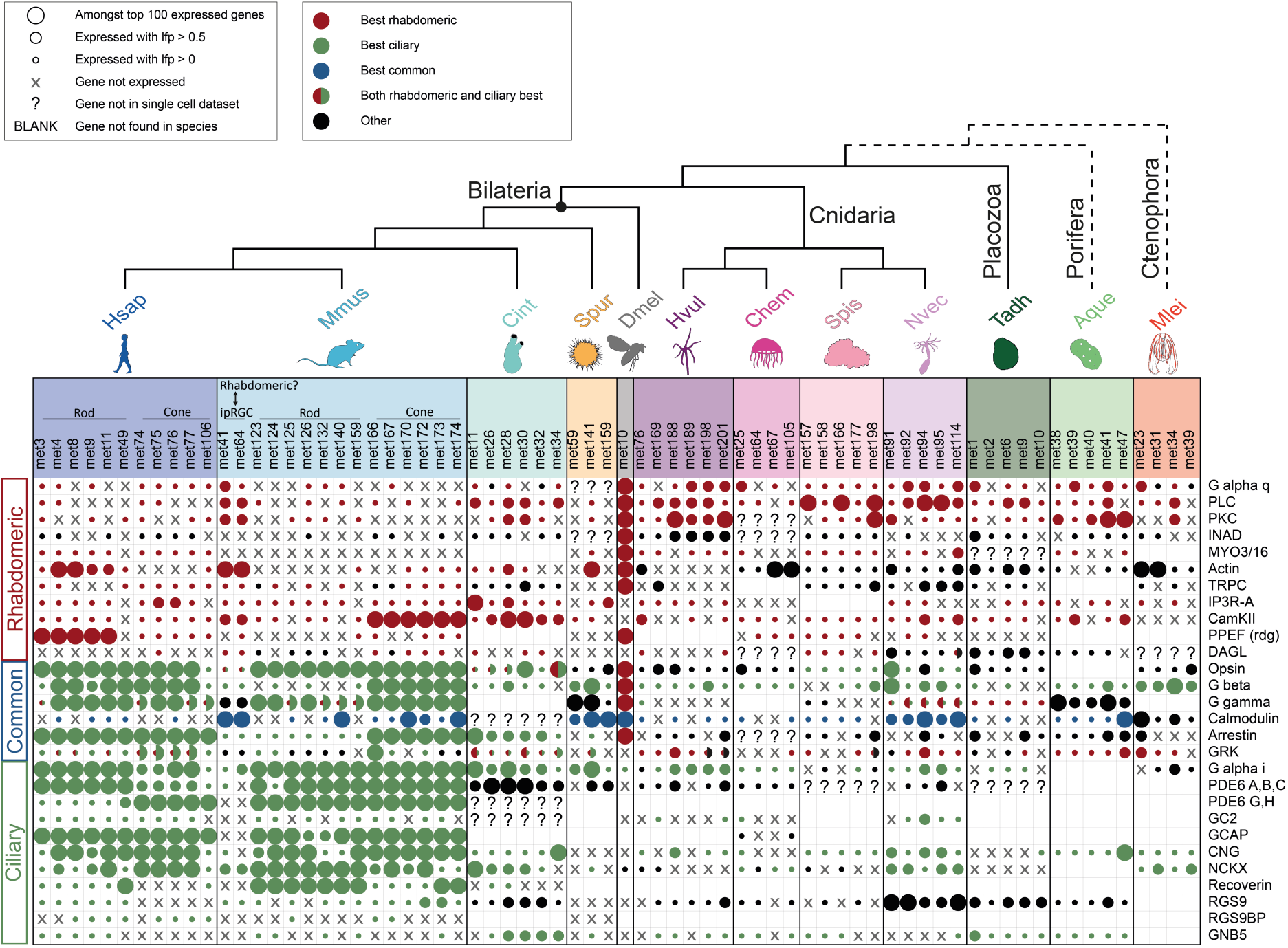
Expression of phototransduction genes in photoreceptor-like cells across animals. The expression of phototransduction genes (listed on the right) is shown for each putative PRC-like metacell across all examined species. Circles represent gene expression, with diameter size proportional to expression levels. A cross denotes that the gene family is not expressed in that metacell, while a question mark indicates that the gene was absent in the single-cell dataset, making expression evaluation impossible. A blank space signifies that the gene family is not present in the genome of that species, rendering expression checks unapplicable. If genes expressed in a metacell belong to a sub-family of interest, circles are colour-coded as red (rhabdomeric), green (ciliary), or blue (common). Black circles indicate that the expressed genes belong to a more distantly related sub-family within the gene family. If both ciliary and rhabdomeric sub-families are co-expressed, the circle is divided into two halves, each with respective colour and size. Human and mouse ciliary PRCs express mainly ciliary type genes, but also some rhabdomeric type ones. Instead, *Drosophila* PRC expresses almost exclusively rhabdomeric type genes. Two candidate ipRGC in mouse display a rhabdomeric-type profile. *Ciona intestinalis* metacells appear to have ciliary-like profiles. Outside of chordates, a large amount of phototransduction genes is either not present in the genome or not detected in the scRNAseq data, and, overall, most species have a mixture of rhabdomeric, and ciliary genes expressed. Species silhouettes were modified from images with CC0 1.0 Universal Public Domain Dedication licences obtained from https://www.phylopic.org/. Abbreviations: Dmel: *Drosophila melanogaster*; Hsap: *Homo sapiens*; Mmu*s*: *Mus musculus*; Cint: *Ciona intestinalis*; Spur: *Strongylocentrotus purpuratus*; Nvec: *Nematostella vectensis*; Spis: *Stylophora pistillata*; Chem: *Clytia hemisphaerica*; Hvul: *Hydra vulgaris*; Tadh: *Trichoplax adhaerens*; Aque: *Amphimedon queenslandica*; Mlei: *Mnemiopsis leidyi*.

**I**nitially, we tested the pipeline in *Drosophila*, humans, and mice, where the composition of PRCs is well known. As expected, mice and human PRC metacells exhibited a strong ciliary profile (**Figure 2**), whereas *Drosophila* PRC showed a strong rhabdomeric profile (**Figure 2**). Additionally, in mice, we identified two metacells with a rhabdomeric profile (**Figure 2**), suggesting that they may represent intrinsically photosensitive retinal ganglion cells (ipRGCs) that have been proposed as homologous to rhabdomeric PRCs (Arendt 2003; Hahn et al. 2023). We then used the same approach to identify putative PRCs in other animals including non-bilaterians, where the presence of PRCs is unclear. For instance, in cnidarians, ciliary and rhadomeric PRCs have been described (Kozmik et al. 2008, Nordström et al. 2003). In ctenophores putative ciliary photoreceptors have been proposed (Horridge 1964). Our results suggest that identifying metacells with a clear ciliary or rhabdomeric transcriptional profile is challenging in non-bilaterian animals. Most metacells express only some components required for the ciliary and rhabdomeric phototransduction cascades. Overall, our findings indicate that, outside of model organisms like mice and *Drosophila* (**Figure 2**), the majority of putative PRC metacells in non-bilaterian animals express a combination of ciliary and rhabdomeric genes, supporting previous observations from Yau and Hardy (Yau and Hardie 2009). We suggest that non-model organisms might use alternative genes for phototransduction.

Integrating phylogenomic and single-cell RNA-seq data allowed the identification of PRCs, in *Drosophila*, mice, and humans and putative PRC-like metacells in non-model organisms, including non-bilaterians. However, the patchy distribution of phototransduction genes or the lack of expression in single-cell data does not clarify the homology of the PRC-like metacells found in non-bilaterian animals. As found in previous works (Burkhardt and Jékely 2021; Musser et al. 2021; Hayakawa et al. 2022), we should be able to identify TFs across animal phyla and identify PRCs. To identify regulatory genes that were highly expressed in each putative PRC we used EggNog (Cantalapiedra et al. 2021) and AnimalTF (Shen et al. 2023). We identified 69 orthogroups of regulatory genes shared across three or more phyla (**Figure 3A and Supplementary Figure S31**), with around 60% classified as TFs (**Figure 3B**), 23% as transcription cofactors that interact with transcription factors but do not directly bind DNA, and ∼17% as other regulatory genes, including RNA polymerases and proteins interacting with the chromatin structure. Overall, we observed that bZIP transcription factors, zinc finger C2H2, and homeoboxes are amongst the most predominant transcription factor families (**Figure 3C**). Furthermore, the variety of DNA-binding-domains suggests a broad spectrum of mechanisms through which transcription can be regulated in photoreceptor-like cells throughout animals (Figure 3C). Among the TFs commonly expressed in three or more phyla, some, like Six6/3, and Tbx2 for ciliary type (Zuber et al. 2003; Alvarez-Delfin et al. 2009; Vopalensky and Kozmik 2009; Valencia et al. 2021) and glass for rhabdomeric (Bernardo-Garcia et al. 2017), are recognised for their role in PRC identity and/or specification. While the exact combination of regulatory genes is not fully conserved, several orthogroups are recurrent throughout animals. The most frequently occurring orthogroups were: ISL, present in metacells spanning 6 species including some cnidarians; TBX/bifid, present in metacells from 5 species including placozoa and ctenophore; and SOX2, present in 5 species including cnidarian, poriferan, and ctenophore metacells (Figure 3A).

**Figure 3.**
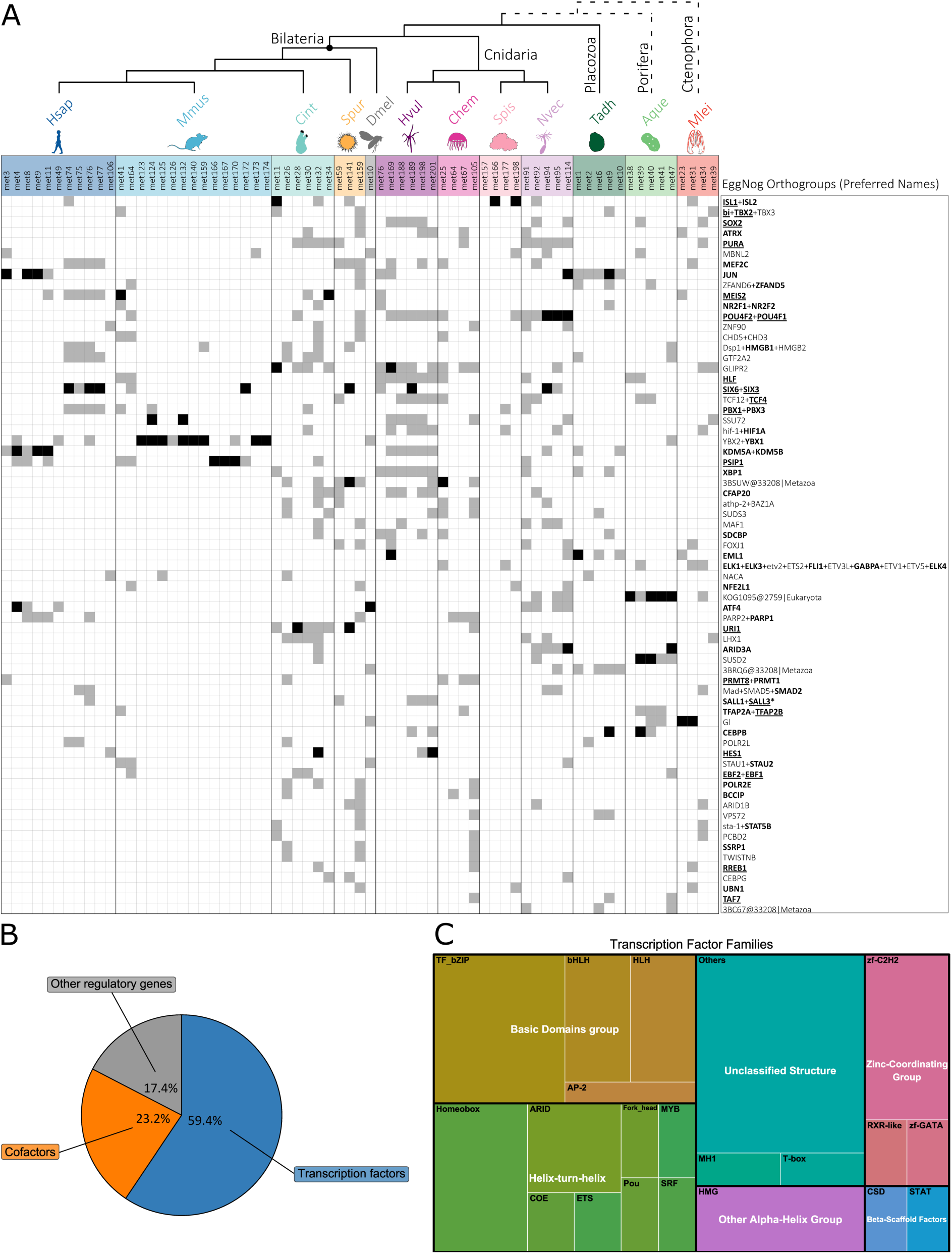
Most common orthogroups of regulatory genes shared across PRC-like metacells throughout animals. (**A**) The 69 orthogroups that are present in 3 or more phyla are listed in order of most frequent (with the hierarchy: present in most phyla, present in most species, present in most metacells). Expression in metacells is indicated with a black square when genes were amongst the top 100 differentially expressed genes of the metacell, while a grey square indicates expression with log-fold enrichment (lfp) > 0.5. Therefore, black squares indicate strong markers for a given metacell, while grey squares indicate that the gene is expressed in the metacell but differential expression level is not as high. Some orthogroups are frequently expressed throughout animals, although their exact combination of co-expression varies in the different species. Orthogroup names derive from the Preferred_names of the respective EggNog orthogroups, where present, or the EggNog orthogroup itself. GeneCards/Flybase were used to characterise the human/*Drosophila* representatives. Genes are highlighted: in bold if there is some evidence of involvement in vision and/or eye/photoreceptor development; in bold and underlined if there is strong evidence of involvement in vision and/or are expressed in the retina, although not necessarily in cones and rods; in bold, underlined and with asterisk if they are specifically expressed in photoreceptor cells. Species silhouettes were modified from images with CC0 1.0 Universal Public Domain Dedication licences obtained from https://www.phylopic.org/. Abbreviations: Dmel: *Drosophila melanogaster*; Hsap: *Homo sapiens*; Mmus: *Mus musculus*; Cint: *Ciona intestinalis*; Spur: *Strongylocentrotus purpuratus*; Nvec: *Nematostella vectensis*; Spis: *Stylophora pistillata*; Chem: *Clytia hemisphaerica*; Hvul: *Hydra vulgaris*; Tadh: *Trichoplax adhaerens*; Aque: *Amphimedon queenslandica*; Mlei: *Mnemiopsis leidyi*. (**B**) The majority of the orthogroups of regulatory genes are transcription factor families (59.4%). Transcription cofactors are also abundant (23.2%). The remaining orthogroups include a mixture of other genes that are involved in transcription, such as polymerases and genes involved in chromatin conformation. (**C**) Treemap of the most abundant families of transcription factors shared across PRC-like metacells, organised by broad groups based on the type of DNA-binding domain. Basic domains and helix-turn-helix are the most abundant.

In cnidarians, our results suggest that although orthologues of several phototransduction genes were missing in the genomes/transcriptomes and/or in the single-cell data, this phylum seems to have a more complete repertoire of phototransduction components compared to other non-bilaterians (Figure 2). *Stylophora pistillata* and *Nematostella vectensis* express ciliary-type opsins in metacells, while *Hydra vulgaris* and *Clytia hemisphaerica* express opsins that are RGR/Go (see Supplementary Figure S7). Together with the ciliary opsins, the presence of other ciliary genes (e.g. CNG, NCKX) suggests a ciliary signature (**Figure 2**), further supported by co-expression with transcription factors such as ISL, HLF, SIX3/6, known to be expressed in retinal progenitor cells in vertebrates and sea urchins (Wu et al. 2015; Valencia et al. 2021; Paganos et al. 2022) (**Figures 3 and 4**). The absence of other ciliary genes, and the presence of several rhabdomeric ones, may reflect the view that cnidarian phototransduction, while sharing some components with the two traditional cascades, also includes cnidaria-specific elements yet to be characterised (Vöcking et al. 2022; Birch et al. 2023). In addition to regulatory genes that cnidarians share with other animals, there are few cases of phyla-specific orthogroups (Supplementary Figure S31). AHR and GATA3 are the most frequently appearing cnidaria-specific orthogroups, with the latter being amongst the most highly expressed (Figure S31 and Table S4).

**Figure 4.**
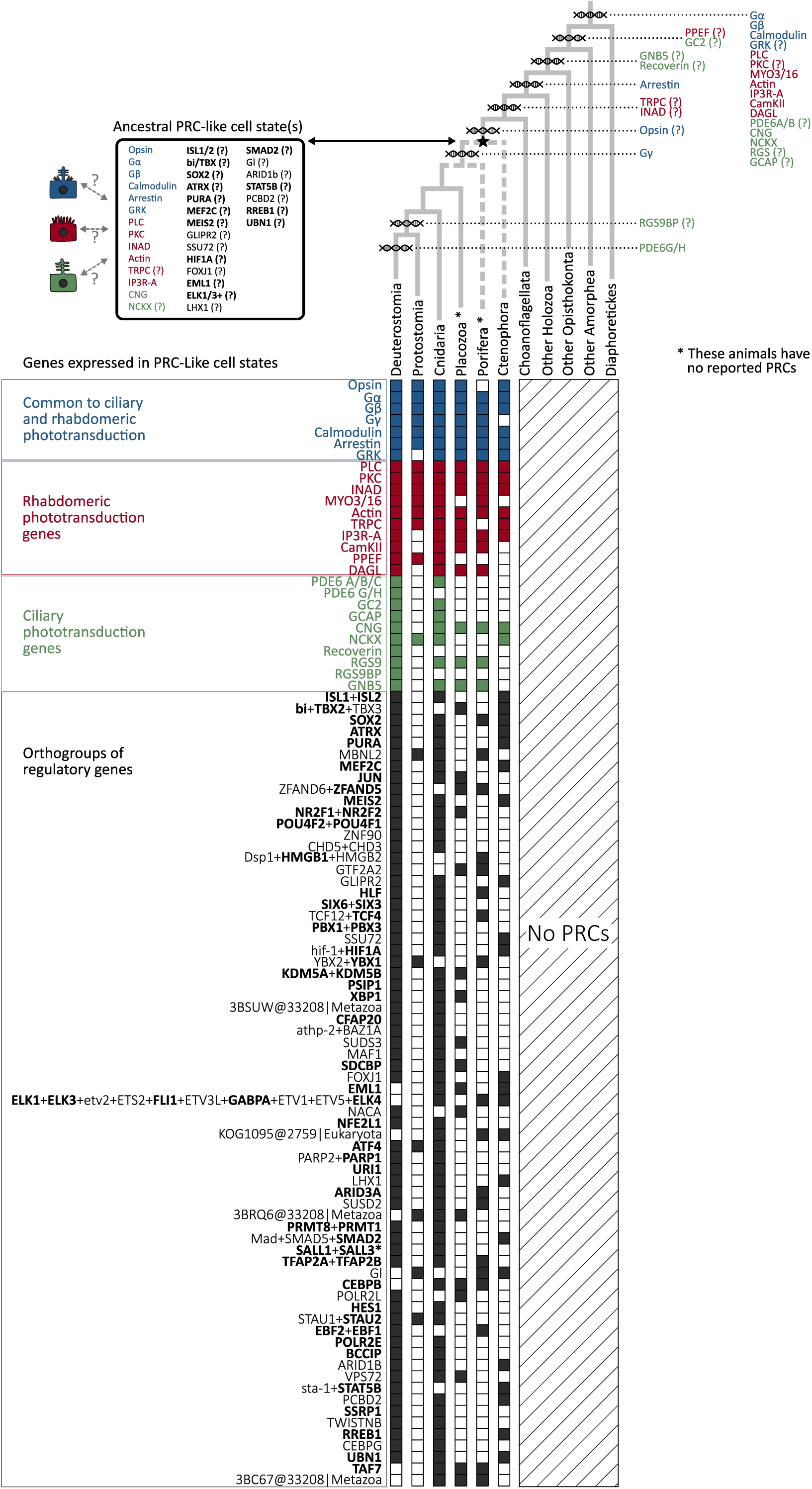
Origin of phototransduction gene families in eukaryotes and expression of phototransduction and regulatory genes in photoreceptor cell-like metacells across animals. The origins of phototransduction gene families are indicated with double-helix symbols on respective branches of the eukaryote species tree (top right). Where there is some uncertainty about the origin of a gene family, such as when the phylogenetic analysis recovered its presence in distantly related eukaryotes, but the gene family has a patchy distribution outside animals/Holozoans, this uncertainty is indicated with a (?) next to the gene family name. A star at the base of animals indicates the origin of an ancestral photoreceptor cell (PRC) type(s). Based on the expression data from PRC-like metacells across animals, the ancestor of modern animals may have had PRC-like(s) with a combination of rhabdomeric, ciliary and common phototransduction genes, as well as several regulatory genes (top left). These may have included up to 6 rhabdomeric phototransduction components, 2 ciliary components and 6 components shared between both pathways, as well as 21 regulatory genes orthogroups, as can be speculated by their presence in PRC-like metacells of ctenophore, the potential sister group of all other animals. Due to the differences in the expression patterns throughout animals, especially regarding regulatory genes orthogroups, a (?) is indicated next to the gene families where there is uneven distribution across animal PRC-likes. In the bottom section of the Figure, the presence/absence of the expression of each family of phototransduction genes and regulatory genes orthogroups in PRC-like metacells is indicated for each animal clade examined, including all non-bilaterian phyla. This presence absence is the result of pooled data from all metacells for a given clade, therefore, a coloured square indicates that the gene family is expressed in at least one PRC-like metacell from at least one species of the clade under consideration.

In the placozoan *Trichoplax adhaerens*, we identified five candidate metacells (**Figure 3 and Table S7**) expressing genes associated with phototransduction, including placopsins (Feuda et al. 2012). Interestingly, from the *Trichoplax* genome, we identified representatives of all eleven rhabdomeric gene families (Figure 1), and these were detected in the single-cell data except the MYO3/16 family (Figure 2). Of these, all except PPEF were expressed in most PRC-like metacells of *Trichoplax*. This contrasts with ciliary genes, of which only a handful were present in the genome, and these were in turn rarely expressed in the PRC-like metacells. Almost all putative PRC metacells in *Trichoplax* expressed the rhabdomeric TF Jun (Bohmann et al. 1994), and one of them the ciliary marker TBX. The most highly expressed regulatory orthogroups in placozoan PRC-like metacells are annotated by EggNog as TSPAN9 and NEUROG1, the first being placozoan-specific and the second appearing also in urochordate metacells (Figure S31 and Table S4).

Ctenophores show a reduced complement of phototransduction genes, however, the analysis of putative PRCs in the ctenophore *M. leydi* identified four candidate PRC-like metacells (Figure 2). Although more rhabdomeric genes are present in the genome compared to ciliary genes (Figure 1), all three ciliary genes (G alpha i/o, CNG and NCKX) available in the genome are expressed in all candidate PRC metacells, in contrast, only a handful of the available rhabdomeric genes are expressed. The presence of putative ciliary PRCs is further supported by the TF expression, including ciliary markers like ISL, TBX, and MEIS2 (Zuber et al. 2003; Alvarez-Delfin et al. 2009; Valencia et al. 2021).

In the sponge *Amphimedon queenslandica* two rhabdomeric genes and most ciliary genes were missing from the genome (Figure 1). Overall, this species, together with the ctenophore, had the fewest phototransduction genes in the genome. In the PRC-like metacells recovered from single-cell analysis, the few ciliary genes found in the genome are all expressed, as are all common genes and most rhabdomeric genes (Figure 2). Unlike other phyla, none of the sponge metacells express well-characterised ciliary or rhabdomeric marker TFs, except for SOX2, where orthologues are expressed in mouse retinal progenitor cells (Lin et al. 2009). Lastly, in both ctenophore and sponge putative PRC-like metacells we found only few species-specific orthogroups and these were neither frequent nor highly expressed. This may either reflect a technical difficulty in detecting species-specific orthogroups in these species or a genuine lack of species-specific orthogroups involved in PRC-like cells of these organisms.

## Discussion

Our study on the evolutionary assembly of vision (summarised in Figure 4) has substantially advanced our understanding on the origins of the visual machinery. Overall, our findings suggest that key components of the phototransduction process evolved from a series of gene duplications in the holozoan and metazoan stem groups that expanded ancient gene families. Additionally, we identified the conservation of PRC regulatory programmes over 600 million years of evolution (Carlisle et al. 2024), suggesting the homology of photoreceptor cells from ctenophores to humans (Figure 4).

Compared to previous studies focused on a subset of genes in few species, our results substantially clarify the duplication pattern of all major phototransduction genes. For example, our findings on opsin are comparable to those obtained by (Fleming et al. 2020); however, they are strongly influenced by the relative phylogenetic positions of ctenophores and sponges, with the ctenophores-first hypothesis supporting a loss of opsin in all sponges. Previous work on CNG suggested that the two subunits, alpha and beta, derived from a duplication at some point before the split of protostomes and deuterostomes (Lamb 2020). Our reconciliation revealed more precisely that this duplication occurred within the ancestor of choanoflagellates and animals (Figure S19). Furthermore, our results clarify the presence of phototransduction genes in ctenophores. A previous study by Schnitzler and collaborators (Schnitzler et al. 2012) using a simple BLAST search identified many ciliary phototransduction genes in *Mnemiopsis leidyi*, in contrast to only a handful of rhabdomeric genes. In contrast, our results suggest a reduced set of classic phototransduction genes in ctenophores. Most likely, this difference is due to methodology, as Schnitzler et al. based homology exclusively on BLAST, while we employed gene-tree to species-tree reconciliation that filtered out genes with dubious homology. Regardless, evidence points towards a ciliary-type PRC in *M. leidyi*, although possibly ctenophores might possess alternative components in their phototransduction cascade.

Our results indicate that ciliary and rhabdomeric phototransduction do not follow the same evolutionary pattern. There are substantial differences in the distribution of the orthologues within animals, where rhabdomeric genes are widely distributed, while ciliary genes are almost uniquely present in vertebrates (Figure 1). Furthermore, our analysis of putative photoreceptor cells suggests that while some shared components were likely utilised early in phototransduction (Figure 2), different animal lineages may have subsequently recruited distinct sets of additional components.

The cross-species comparison of putative PRCs identified several TFs shared between phyla, including key regulators of PRC cell identity such as TBX and ISL (Figure 3). Remarkably, some of these TFs are conserved in non-bilaterians. In cnidarians, where previous evidence suggests the presence of ciliary PRCs (Kozmik et al. 2008) as well as rhabdomeric PRCs in larvae (Nordström et al. 2003), we find a high number of TFs (e.g., ISL, SOX2, SIX3/6) shared with deuterostomes (Figure 4), confirming the presence of putative ciliary PRCs. PRCs with a ciliary morphology have been described in the ctenophore *Pleurobrachia pileus* (Horridge 1964), and although obtained in a different species, our results corroborate this, suggesting that putative PRC cells express the ciliary TFs markers TBX and ISL.

Overall, our findings, like those of (Hayakawa et al. 2022) and (Burkhardt and Jékely 2021), indicate that some neuronal cell types seem conserved in ctenophores despite the dramatic differences in their nervous system morphology (Sachkova et al. 2021; Burkhardt et al. 2023). Conversely, placozoans have a very simple body plan in which only a handful of cell types have been described morphologically (Smith et al. 2014), although molecular studies have uncovered a broader diversity of cell types (Sebé-Pedrós, Chomsky, et al. 2018; Varoqueaux et al. 2018). While *Trichoplax* seems to have at least some simple response to light (Heyland et al. 2014), there is no morphological evidence of the presence of photoreceptor cells. Our results suggest that putative metacells in *Trichoplax* express the rhabdomeric TF Jun (Bohmann et al. 1994) and one of them the ciliary marker TBX in combination with placopsins (Figure 3). Yet, the functions of these putative PRCs and their potential role in light response remain unclear.

Although sponges lack opsins and neurons, they are known to be receptive to light (Leys and Degnan 2001; Maldonado et al. 2003; Elliott and Leys 2004; Wong et al. 2022). It has been proposed that sponges may utilise light-sensitive cryptochromes (Rivera et al. 2012; Müller et al. 2013) or other GPCRs such as glutamate receptors (Wong et al. 2022) for photoreception. In *Amphimedon queenslandica*, two rhabdomeric phototransduction genes have been implicated in the phototactic behaviour of larvae (Wong et al. 2022). Our results suggest that phototransduction genes are expressed in the same metacells together with Sox2 TF genes, usually expressed in retinal progenitor cells in mice (Lin et al. 2009), suggesting the existence of a phototransduction pathway and some elements of the photoreceptor regulatory programme in these animals.

Our current study is subject to certain limitations. First, some species’ genomes are incomplete, which might affect our pattern of duplications and losses. However, using multiple species per phylum partially mitigates this issue. Second, cross-phyla comparison of single-cell RNA-seq data is challenging and exacerbated by the use of different technologies, resulting in the detection of varying numbers of genes per cell, affecting our ability to detect and compare the expression of orthologous genes. Additionally, annotation tools such as EggNog may produce less accurate results for non-bilaterian species and this needs to be considered when interpreting results. Finally, defining homology of cell types is challenging. While it is evident that some TFs must be shared, their exact number and combination is unclear. To overcome this challenge, it will be fundamental understand the TFs that control neuronal identity. In the case of PRCs, the TFs that controls the expressions of opsins and phototransduction genes. Despite these limitations, the duplication patterns of phototransduction genes and the conservation of TF expression in putative PRCs suggest a scenario for the origin of vision where the duplication of pre-existing families in the metazoan stem group gave rise to key components of the phototransduction cascade. Subsequently, these newly evolved genes underwent regulatory control by key TF families, conserved across most animals. Starting from these ancestral cells, these regulatory programmes underwent lineage-specific diversification, giving rise to the diversity of PRCs observed in modern-day metazoans.

## Materials and Methods

### Species Selection and Proteome Assessment

We selected 86 representative species of eukaryotes based on proteome completeness and taxonomic distribution (Supplementary Table S2). Our focus was on sister taxa of Metazoa, including 8 choanoflagellates and 5 other holozoans, as well as 25 non-bilaterian Metazoa species. This selection was motivated by the need to understand the origins of functional visual processes at an early stage of animal evolution. Proteome completeness was assessed using BUSCO (v4.0.6) (Simão et al. 2015; Waterhouse et al. 2018) with the eukaryota_odb10 database comprising 255 BUSCO genes. Supplementary Table S2 provides BUSCO scores for each proteome.

### Species Phylogeny Construction

To estimate the species phylogeny, we utilised single-copy BUSCO orthologs. BUSCO genes were extracted and aligned using MAFFT v7.470 (--auto) (Katoh et al. 2002; Katoh and Standley 2013). Alignments were then trimmed with Trimal v1.4 (-automated1) (Capella-Gutiérrez et al. 2009). Trimmed alignments of all BUSCO genes were concatenated into a super-matrix using FASconCAT v1.11 (Kück and Meusemann 2010). This super-matrix served as input for species tree construction with IQTREE v2.0.6 (Hoang et al. 2018; Minh et al. 2020), using Model Finder (Kalyaanamoorthy et al. 2017) to determine the best-fitting model (LG+F+R10). The species tree positioned Ctenophores as the sister group to all other metazoans (Whelan et al. 2017; Schultz et al. 2023). An alternative topology, with sponges as the sister group to all other animals (Feuda et al. 2017), was obtained by manually swapping branches using Mesquite v3.6.1 (Maddison and Maddison 2008). Both topologies were used for the reconciliation and are available on FigShare (see data availability section).

### Identification of Phototransduction Gene Homologs

To identify the homologs of the phototransduction genes, we used KEGG maps. BLAST queries for a total of 28 gene families were identified primarily based on the KEGG maps ko04745 (rhabdomeric) and ko04744 (ciliary) (Kanehisa et al. 2021). Additionally, the RGS9BP and GNB5 subunits of the RGS9 complex were included based on updated references of vertebrate phototransduction (Lamb et al. 2018) (see Supplementary Figure S1 and Supplementary Table S1). BLASTP searches were conducted (e-value cut-off of 1e-5) for each query versus the species database. Potential duplicates were removed with cd-hit (Li et al. 2001; Fu et al. 2012) with an identity threshold of 100%. Putative homologs were used for another BLASTP versus the SwissProt database (Poux et al. 2017). Sequences were kept if the gene family of interest was within the top five hits, parsed using gene family-specific keywords (**see Table S5**). The first level of filtering was based on similarity. A second round of filtering was conducted based on the presence of gene family-specific protein motifs. The dataset was scanned with InterProScan (Quevillon et al. 2005; Jones et al. 2014) and sequences were retained only if they contained the characteristic combination of motifs (**Table S5**). To provide an annotation to the final collections of sequences, we used the top hit from BLASTP versus SwissProt.

### Phylogenetic Analysis

Gene trees were constructed for each gene family following a standard pipeline. Sequences were aligned with MAFFT (--auto) (Katoh et al. 2002; Katoh and Standley 2013) and trimmed to eliminate columns with more than 70% gaps using Trimal (-gt 0.3) (Capella-Gutiérrez et al. 2009). Tree construction was carried out in IQTREE2 using the best-fitting models according to BIC (**see Table S6**). Branch supports were assessed with 1000 replicates for the Shimodaira–Hasegawa-like approximate likelihood ratio test (-alrt) (Guindon et al. 2010), the Bayesian-like transformation of aLRT (-abayes) (Anisimova et al. 2011), and with 1000 replicates for ultrafast bootstrap approximation (-B) (Minh et al. 2013; Hoang et al. 2018). To obtain fully bifurcating gene trees necessary as inputs for the gene tree to species tree reconciliations, any polytomy in the gene trees was randomly resolved with ETE3 (Huerta-Cepas et al. 2016).

### Gene Tree to Species Tree Reconciliation

The gene trees were used as starting trees for gene tree-to-species tree reconciliations using GeneRax (v1.2.3) (Morel et al. 2020). The model used to compute the reconciliation was set to account for duplication and loss but not transfer events. Both alternative species trees (ctenophore-first and sponge-first) were tested. The number of duplications and losses for each gene family were compared between scenarios **(see Table S3**).

The resulting reconciled trees were manually examined to trace the evolution of the genes of interest. The *D. melanogaster* and *H. sapiens* genes known to function in phototransduction were used to identify the orthogroups of interest and the duplication and loss events that characterized their lineages. Other subgroups within the gene families and their relationship with the orthogroups of interest were also identified.

By tracing the presence of the orthologs identified from reconciliations throughout all the species examined, we compiled a list of candidate marker genes for phototransduction in non-model organisms. Where the orthologs of interest were not present, paralogs were used as potential markers. These marker genes were used for identifying candidate photoreceptor cell types in non-model organisms, including several non-bilaterians, for which single-cell RNA sequencing data was available.

### Identification of Putative Photoreceptor Cell Types from Single-Cell RNA-Sequencing Data

#### Species Datasets

To obtain a sample of photoreceptor cell diversity throughout Metazoa, we focused the single-cell analysis on twelve species based on scRNA-seq data availability and phylogenetic representation. *Drosophila melanogaster* (Özel et al. 2021) served as an example for rhabdomeric-type PRCs, while *Homo sapiens* (Lukowski et al. 2019) and *Mus musculus* (Macosko et al. 2015) were representative for ciliary-type PRCs. Two additional deuterostomes, the urochordate *Ciona intestinalis* (Sharma et al. 2019) and the sea urchin *Strongylocentrotus purpuratus* (Paganos et al. 2021), served as bridge species between vertebrate PRCs and protostome PRCs, as represented by *Drosophila*.

Non-bilaterian animals of particular interest included four cnidarian species: *Hydra vulgaris* (Siebert et al. 2019), *Clytia hemisphaerica* (Chari et al. 2021), *Stylophora pistillata* (Levy et al. 2021), and *Nematostella vectensis* (Sebé-Pedrós, Saudemont, et al. 2018). We also included the placozoan *Trichoplax adhaerens* (Sebé-Pedrós, Chomsky, et al. 2018), the sponge *Amphimedon queenslandica* (Sebé-Pedrós, Chomsky, et al. 2018), and the ctenophore *Mnemiopsis leidyi* (Sebé-Pedrós, Chomsky, et al. 2018). The details of these scRNA-seq datasets are summarized in Figure S30.

### MetaCell Pipeline for Clustering Cells

Molecular count matrices for each species were used as input for an established pipeline using the MetaCell v0.3.6 (Baran et al. 2019) R package, as described on the MetaCell GitHub (https://tanaylab.github.io/metacell/articles/d-amphimedon.html).

We developed a pipeline to identify metacells that most likely were PRCs based on the expression of combinations of phototransduction genes. First, we filtered out metacells with opsin log fold enrichment (lfp) below 0.2, as opsin is a strong marker for photoreceptor cells. The exception was *Amphimedon queenslandica*, as sponges do not possess opsins (Feuda et al. 2012). In this case we used other phototransduction genes as markers. Next, we assessed the level of phototransduction gene expression in the metacells, checking both the percentage of phototransduction genes co-expressed in the same metacell and their level of expression. Specifically, we calculated the percentage of phototransduction genes expressed and their average lfp for all genes, common genes, rhabdomeric genes, and ciliary genes. To classify metacells into the best PRC candidates, we considered the following evidence: 1) lfp of the highest expressed opsin in the metacell, 2) average lfp of all phototransduction genes, 3) average lfp of common phototransduction genes, 4) average lfp of either ciliary or rhabdomeric genes (whichever is highest), 5) the highest percentage of all phototransduction genes, 6) the highest percentage of common phototransduction genes, and 7) the highest percentage of either ciliary or rhabdomeric phototransduction genes (whichever is highest). We ranked the metacells from best to worst for each category and summed the ranking values to obtain a final ranking. We retained up to the top 5 metacells as the best candidate PRCs for further analyses. Separate rankings were made for rod and cones in human and mouse (**see Table S7**).

### Exploring the Regulatory Toolkit of Candidate PRCs and Comparison Across Species

After identifying PRC-like metacells based on phototransduction gene expression, we further characterized the genetic profile of these candidate PRCs, focusing on regulatory genes such as transcription factors, which define cell identity (Arendt et al. 2016). For all candidate PRCs, we collected the top 100 most highly expressed genes as additional markers and all genes with a lfp above 0.5, representing genes mildly overexpressed in the given metacell.

We used two tools to identify genes involved in transcription. First, we annotated all collected genes with Eggnog mapper (Cantalapiedra et al. 2021), retaining only genes involved in transcription regulation (K category). Additionally, we scanned sequences for Pfam profiles of known TFs (**see Table S8**). Combining these approaches, we compiled a list of transcription factors and genes involved in transcription for all metacells. For cross-species comparison, we used the Eggnog Orthogroup (Eggnog_OG) of the genes preferably at Metazoa level, or the most stringent level available (e.g., Opisthokonta or Eukarya).

### Uncovering Common Regulatory Genes in PRC-Like Metacells

We mapped the presence/absence of regulatory genes orthogroups across species and ordered them by frequency (**Figure 3A and Figure S31**). To classify the orthogroups into transcription factors and other regulatory genes (e.g. transcription cofactors), we performed BLASTP against the Animal Transcription Factor Database (ATFDB version 4) (Shen et al. 2023) (Figure 3B). We further categorized transcription factors into families and broader groups based on the DNA-binding domain (**Figure 3C**).

## Data Availability

Supplementary material and raw output files are available on FigShare: https://doi.org/10.25392/leicester.data.c.7284370. The scripts used for the metacell analyses are also available at the GitHub repository: https://github.com/AAleotti/EvoPhototrPRCs.

## Acknowledgements

This work is supported by a University Research Fellowship (UF160226 and URF\R\221011) to R. Feuda. A. Aleotti is supported by a Research Grant from The Royal Society to R Feuda (RGF\R1\181012). This research used the ALICE High-Performance Computing facility at the University of Leicester. The authors are grateful to Maryam Ghaffari Saadat and Julien Devilliers for their availability to help with coding and automating steps used in the methodologies described in this paper.

