## Supplementary material for "A Pre-Bilaterian Origin of Phototransduction Genes and Photoreceptor Cells": TableS1

| **Common** | **Component**  **(Gene Family)** | ***Drosophila melanogaster*** | | ***Homo sapiens*** | |
| --- | --- | --- | --- | --- | --- |
|  |  | **Gene Name(s)** | **Protein Name(s)** | **Gene Name(s)** | **Protein Name(s)** |
|  | Opsin | ninaE | Opsin Rh1 | OPN1LW | Long-wave-sensitive opsin 1 |
|  |  | Rh 2 to 7 | Opsins Rh2 to 7 | OPN1SW | Short-wave-sensitive opsin 1 |
|  |  |  |  | OPN1MW | Medium-wave-sensitive opsin 1 |
|  |  |  |  | RHO | Rhodopsin |
|  | G beta | Gbeta76C | Gbeta76C (or Gbe) | GNB 1 to 4 | G protein subunit beta 1 to 4 |
|  | G gamma | Ggamma30A | Ggamma(e) | GNGT1 | G protein G(T) subunit gamma-T1 |
|  |  |  |  | GNGT2 | G protein G(T) subunit gamma-T2 |
|  | Calmodulin | Cam | Calmodulin (or CaM) | CALM 1 to 3 | Calmodulin 1 to 3 |
|  | Arrestin | Arr1 | Phosrestin-2 (or Arrestin-1) | ARR3 | Arrestin-C |
|  |  | Arr2 | Phosrestin-1 (or Arrestin-2) | SAG | S-arrestin |
|  | GRK | Gprk1 | GPCR kinase 1 | GRK1 | Rhodopsin kinase GRK1 |
|  |  |  |  | GRK7 | Rhodopsin kinase GRK7 |
| **Rhabdomeric** | **Component**  **(Gene Family)** | ***Drosophila melanogaster*** | | | |
|  |  | **Gene Name(s)** | **Protein Name(s)** | | |
|  | G alpha q | Galphaq | G protein alpha q subunit | | |
|  | PLC | norpA | Phosphoinositide phospholipase C-beta | | |
|  | PKC | inaC | Protein kinase C, eye isozyme (or Eye-PKC) | | |
|  | INAD | inaD | Inactivation-no-after-potential D protein | | |
|  | MYO3 | ninaC | Neither inactivation nor afterpotential protein C | | |
|  | Actin | Act5C | Actin-5C | | |
|  | TRP C | trp | Transient receptor potential protein | | |
|  |  | trpl | Transient-receptor-potential-like protein | | |
|  | IP3R-A | Itpr | Inositol 1,4,5-trisphosphate receptor (or IP3R) | | |
|  | CamKII | CaMKII | Calcium/calmodulin-dependent protein kinase | | |
|  | DAGL | inaE | Inactivation no afterpotential E | | |
|  | PPEF | rdgC | Serine/threonine-protein phosphatase rdgC (or Retinal degeneration C protein) | | |
| **Ciliary** | **Component**  **(Gene Family)** | ***Homo sapiens*** | | | |
|  |  | **Gene Name(s)** | **Protein Name(s)** | | |
|  | G alpha i | GNAT1 | G protein G(t) subunit alpha 1 | | |
|  |  | GNAT2 | G protein G(t) subunit alpha 2 | | |
|  | PDE6 A/B/C | PDE6A | Rod cGMP-specific 3',5'-cyclic phosphodiesterase subunit alpha | | |
|  |  | PDE6B | Rod cGMP-specific 3',5'-cyclic phosphodiesterase subunit beta | | |
|  |  | PDE6C | Cone cGMP-specific 3',5'-cyclic phosphodiesterase subunit alpha' | | |
|  | PDE6 G/H | PDE6G | Retinal rod rhodopsin-sensitive cGMP 3',5'-cyclic phosphodiesterase subunit gamma | | |
|  |  | PDE6H | Retinal cone rhodopsin-sensitive cGMP 3',5'-cyclic phosphodiesterase subunit gamma | | |
|  | GC2 | GUCY2D | Retinal guanylyl cyclase 1 (or GC2D) | | |
|  |  | GUCY2F | Retinal guanylyl cyclase 2 (or GC2F) | | |
|  | GCAP | GUCA1A | Guanylyl cyclase-activating protein 1 (GCAP1) | | |
|  |  | GUCA1B | Guanylyl cyclase-activating protein 2 (GCAP2) | | |
|  |  | GUCA1C | Guanylyl cyclase-activating protein 3 (GCAP3) | | |
|  | CNG | CNGA 1 to 4 | cGMP-gated cation channel alpha 1 to 4 | | |
|  |  | CNGB 1 and 3 | Cyclic nucleotide-gated cation channel beta 1 and 3 | | |
|  | NCKX | SLC24A 1, 2 and 4 | Sodium/potassium/calcium exchanger 1, 2 and 4 (or NCKX1,2 and 4) | | |
|  | Recoverin | RCVRN | Recoverin | | |
|  | RGS9 | RGS9 | Regulator of G-protein signaling 9 | | |
|  | RGS9BP | RGS9BP | Regulator of G-protein signaling 9-binding protein (RGS9BP or RGBP) or RGS9-anchoring protein (R9AP) | | |
|  | GNB5 | GNB5 | G protein subunit beta 5 | | |
