## Supplementary material for "A Pre-Bilaterian Origin of Phototransduction Genes and Photoreceptor Cells": Caption

Supplementary Figure Legends

Supplementary Figure S1. Schematics of rhabdomeric and ciliary phototransduction pathways. (A) Rhabdomeric phototransduction in *Drosophila melanogaster* occurs in the microvilli of the rhabdomere, a structure within the cell body of the photoreceptor cell. The opsin interacts with a G alpha q that activates phospholipase C (PLC) initiating a phosphoinositide cascade that culminates in depolarisation of the photoreceptor cell. (B) Ciliary phototransduction in *Homo sapiens* occurs in a specialised cilium of the photoreceptor cell. The opsin activates the G alpha of transducin that in turn activates phosphodiesterase 6 with consequent cascade that causes the hyperpolarization of the photoreceptor cell. In rod photoreceptors, the opsin and the other membrane proteins, with the exception of the ion channels, are in the membrane of the disk as depicted here. In cone photoreceptors, whilst the components are the same, all membrane components are in the cell membrane (not depicted here). The pathways are based primarily on the Kegg maps ko04745 (rhabdomeric) and ko04744 (ciliary). Protein components are coloured in red (rhabdomeric pathway) or green (ciliary pathway), ions and other non-protein molecules are represented by small grey circles. Lines between components indicate physical interaction, normal arrows between components indicate activation, normal arrows through channels indicate passage of ions, inhibitory arrows indicate inactivation, dotted arrows indicate movement/ transition towards, +p indicates phosphorylation, -p indicates de-phosphorylation, ? indicates unclear mechanism.

**Supplementary Figure S2.** **Annotated arrestin protein family tree reconciled with a ctenophore-first species tree.** The tree was inferred using GeneRax and the RecPhyloxml output file was annotated. In this format, the gene tree is represented within the species tree. Major species groups are indicated on the right of the tree. Sub-lineages within the gene family are represented by coloured lines. Splits at species nodes represent speciation events. Gene duplications and losses are indicated with a square and a cross, respectively. A half-red, half-green star represents the duplication of origin of the two lineages used in rhabdomeric and ciliary phototransduction. Other important duplication events are annotated throughout the tree.

**Supplementary Figure S3.** **Annotated calmodulin protein family tree reconciled with a ctenophore-first species tree.** The tree was inferred using GeneRax and the RecPhyloxml output file was annotated. In this format, the gene tree is represented within the species tree. Major species groups are indicated on the right of the tree. Sub-lineages within the gene family are represented by coloured lines. Splits at species nodes represent speciation events. Gene duplications and losses are indicated with a square and a cross, respectively. Both rhabdomeric and ciliary calmodulins are within the lineage coloured in blue. A half-blue, half-light blue star represents the duplication that gives rise to calmodulin, and its paralog lineage, calmodulin-like 3. Other important duplication events are annotated throughout the tree.

**Supplementary Figure S4.** **Annotated G beta protein family tree reconciled with a ctenophore-first species tree.** The tree was inferred using GeneRax and the RecPhyloxml output file was annotated. In this format, the gene tree is represented within the species tree. Major species groups are indicated on the right of the tree. Sub-lineages within the gene family are represented by coloured lines. Splits at species nodes represent speciation events. Gene duplications and losses are indicated with a square and a cross, respectively. The lineage of rhabdomeric (red) and ciliary (green) G beta subgroups, are not direct paralogs. A half-red, half-green star represents the duplication that gave rise to rhabdomeric and ciliary G beta as well as other related lineages. Other important duplication events are annotated throughout the tree.

**Supplementary Figure S5. Annotated G gamma protein family tree reconciled with a ctenophore-first species tree.** The tree was inferred using GeneRax and the RecPhyloxml output file was annotated. In this format, the gene tree is represented within the species tree. Major species groups are indicated on the right of the tree. Sub-lineages within the gene family are represented by coloured lines. Splits at species nodes represent speciation events. Gene duplications and losses are indicated with a square and a cross, respectively. A half-red, half-green star represents the duplication of origin of the two lineages used in rhabdomeric and ciliary phototransduction. Other important duplication events are annotated throughout the tree.

**Supplementary Figure S6. Annotated GRK protein family tree reconciled with a ctenophore-first species tree.** The tree was inferred using GeneRax and the RecPhyloxml output file was annotated. In this format, the gene tree is represented within the species tree. Major species groups are indicated on the right of the tree. Sub-lineages within the gene family are represented by coloured lines. Splits at species nodes represent speciation events. Gene duplications and losses are indicated with a square and a cross, respectively. The lineage of rhabdomeric (red) and ciliary (green) GRKs are not direct paralogs. A half-green, half-blue star represents the duplication of origin between the ciliary GRKs lineage and its direct paralogous lineage. A half-red, half-pink star indicates the duplication between the rhabdomeric GRKs and all other animal GRKs. Other important duplication events are annotated throughout the tree.

**Supplementary Figure S7. Annotated opsin protein family tree reconciled with a ctenophore-first species tree.** The tree was inferred using GeneRax and the RecPhyloxml output file was annotated. In this format, the gene tree is represented within the species tree. Major species groups are indicated on the right of the tree. Sub-lineages within the gene family are represented by coloured lines. Splits at species nodes represent speciation events. Gene duplications and losses are indicated with a square and a cross, respectively. A half-red, half-green star represents the duplication that gives rise to ciliary opsins, rhabdomeric opsins, and RGR/Go opsins. Other important duplication events are annotated throughout the tree.

**Supplementary Figure S8. Annotated actin protein family tree reconciled with a ctenophore-first species tree.** The tree was inferred using GeneRax and the RecPhyloxml output file was annotated. In this format, the gene tree is represented within the species tree. Major species groups are indicated on the right of the tree. Sub-lineages within the gene family are represented by coloured lines. Splits at species nodes represent speciation events. Gene duplications and losses are indicated with a square and a cross, respectively. A multi-pointed red star represents the duplication that gave rise to the lineage of muscle and cytoplasmic actins, which includes the rhabdomeric phototransduction component. Other important duplication events are annotated throughout the tree.

**Supplementary Figure S9. Annotated CamKII protein family tree reconciled with a ctenophore-first species tree.** The tree was inferred using GeneRax and the RecPhyloxml output file was annotated. In this format, the gene tree is represented within the species tree. Major species groups are indicated on the right of the tree. Sub-lineages within the gene family are represented by coloured lines. Splits at species nodes represent speciation events. Gene duplications and losses are indicated with a square and a cross, respectively. A multi-pointed red star represents the duplication that gave rise to the lineage of CamKII, which includes the rhabdomeric phototransduction component. Other important duplication events are annotated throughout the tree.

**Supplementary Figure S10. Annotated DAGL protein family tree reconciled with a ctenophore-first species tree.** The tree was inferred using GeneRax and the RecPhyloxml output file was annotated. In this format, the gene tree is represented within the species tree. Major species groups are indicated on the right of the tree. Sub-lineages within the gene family are represented by coloured lines. Splits at species nodes represent speciation events. Gene duplications and losses are indicated with a square and a cross, respectively. A half-red, half-light red star represents the duplication that gave rise to the lineage of DAGL alpha and beta, which includes the rhabdomeric phototransduction component. Other important duplication events are annotated throughout the tree.

**Supplementary Figure S11. Annotated G alpha q protein family tree reconciled with a ctenophore-first species tree.** The tree was inferred using GeneRax and the RecPhyloxml output file was annotated. In this format, the gene tree is represented within the species tree. Major species groups are indicated on the right of the tree. Sub-lineages within the gene family are represented by coloured lines. Splits at species nodes represent speciation events. Gene duplications and losses are indicated with a square and a cross, respectively. A multi-pointed red star represents the duplication of origin of GNAQ and related G alpha proteins. Other important duplication events are annotated throughout the tree.

**Supplementary Figure S12. Annotated INAD protein family tree reconciled with a ctenophore-first species tree.** The tree was inferred using GeneRax and the RecPhyloxml output file was annotated. In this format, the gene tree is represented within the species tree. Major species groups are indicated on the right of the tree. Sub-lineages within the gene family are represented by coloured lines. Splits at species nodes represent speciation events. Gene duplications and losses are indicated with a square and a cross, respectively. A multi-pointed red star represents the duplication of origin of the rhabdomeric phototransduction component. Other important duplication events are annotated throughout the tree.

**Supplementary Figure S13. Annotated IP3R-A protein family tree reconciled with a ctenophore-first species tree.** The tree was inferred using GeneRax and the RecPhyloxml output file was annotated. In this format, the gene tree is represented within the species tree. Major species groups are indicated on the right of the tree. Sub-lineages within the gene family are represented by coloured lines. Splits at species nodes represent speciation events. Gene duplications and losses are indicated with a square and a cross, respectively. A half-red, half-light red star represents the duplication of origin of IP3R-A, including the rhabdomeric phototransduction component, and its paralog. Other important duplication events are annotated throughout the tree.

**Supplementary Figure S14. Annotated MYO3/MYO16 protein family tree reconciled with a ctenophore-first species tree.** The tree was inferred using GeneRax and the RecPhyloxml output file was annotated. In this format, the gene tree is represented within the species tree. Major species groups are indicated on the right of the tree. Sub-lineages within the gene family are represented by coloured lines. Splits at species nodes represent speciation events. Gene duplications and losses are indicated with a square and a cross, respectively. A half-red, half-light red star represents the duplication that gives rise to MYO3, which includes the *Drosophila* ninaC, and MYO16. Other important duplication events are annotated throughout the tree.

**Supplementary Figure S15. Annotated PKC protein family tree reconciled with a ctenophore-first species tree.** The tree was inferred using GeneRax and the RecPhyloxml output file was annotated. In this format, the gene tree is represented within the species tree. Major species groups are indicated on the right of the tree. Sub-lineages within the gene family are represented by coloured lines. Splits at species nodes represent speciation events. Gene duplications and losses are indicated with a square and a cross, respectively. A half-red, half-light red star represents the duplication of origin of classical PKCs, including the rhabdomeric phototransduction component, and novel PKCs. Other important duplication events are annotated throughout the tree.

**Supplementary Figure S16. Annotated PLC protein family tree reconciled with a ctenophore-first species tree.** The tree was inferred using GeneRax and the RecPhyloxml output file was annotated. In this format, the gene tree is represented within the species tree. Major species groups are indicated on the right of the tree. Sub-lineages within the gene family are represented by coloured lines. Splits at species nodes represent speciation events. Gene duplications and losses are indicated with a square and a cross, respectively. A multi-pointed red star represents the duplication of origin of PLC beta, which includes the rhabdomeric phototransduction component. Other important duplication events are annotated throughout the tree.

**Supplementary Figure S17. Annotated PPEF protein family tree reconciled with a ctenophore-first species tree.** The tree was inferred using GeneRax and the RecPhyloxml output file was annotated. In this format, the gene tree is represented within the species tree. Major species groups are indicated on the right of the tree. Sub-lineages within the gene family are represented by coloured lines. Splits at species nodes represent speciation events. Gene duplications and losses are indicated with a square and a cross, respectively. A multi-pointed red star represents the duplication of origin of PPEF, including the rhabdomeric phototransduction component. Other important duplication events are annotated throughout the tree.

**Supplementary Figure S18. Annotated TRPC protein family tree reconciled with a ctenophore-first species tree.** The tree was inferred using GeneRax and the RecPhyloxml output file was annotated. In this format, the gene tree is represented within the species tree. Major species groups are indicated on the right of the tree. Sub-lineages within the gene family are represented by coloured lines. Splits at species nodes represent speciation events. Gene duplications and losses are indicated with a square and a cross, respectively. A multi-pointed red star represents the duplication of origin of TRPC 1/4/5 lineage, which includes the *Drosophila* trp and trpl used in rhabdomeric phototransduction. Other important duplication events are annotated throughout the tree.

**Supplementary Figure S19. Annotated CNG protein family tree reconciled with a ctenophore-first species tree.** The tree was inferred using GeneRax and the RecPhyloxml output file was annotated. In this format, the gene tree is represented within the species tree. Major species groups are indicated on the right of the tree. Sub-lineages within the gene family are represented by coloured lines. Splits at species nodes represent speciation events. Gene duplications and losses are indicated with a square and a cross, respectively. A half-green, half-dark green star represents the duplication of origin of the lineage of CNGA and CNGB, which includes the ciliary phototransduction components. Other important duplication events are annotated throughout the tree.

**Supplementary Figure S20. Annotated G alpha i/o protein family tree reconciled with a ctenophore-first species tree.** The tree was inferred using GeneRax and the RecPhyloxml output file was annotated. In this format, the gene tree is represented within the species tree. Major species groups are indicated on the right of the tree. Sub-lineages within the gene family are represented by coloured lines. Splits at species nodes represent speciation events. Gene duplications and losses are indicated with a square and a cross, respectively. A multi-pointed green star represents the duplication of origin of the lineage that includes GNAT, involved in ciliary phototransduction, and GNAI/Z. Other important duplication events are annotated throughout the tree.

**Supplementary Figure S21. Annotated GC2 protein family tree reconciled with a ctenophore-first species tree.** The tree was inferred using GeneRax and the RecPhyloxml output file was annotated. In this format, the gene tree is represented within the species tree. Major species groups are indicated on the right of the tree. Sub-lineages within the gene family are represented by coloured lines. Splits at species nodes represent speciation events. Gene duplications and losses are indicated with a square and a cross, respectively. A multi-pointed green star represents the duplication of origin of GC2 D/E/F/G, that includes the ciliary phototransduction components. Other important duplication events are annotated throughout the tree.

**Supplementary Figure S22. Annotated GCAP protein family tree reconciled with a ctenophore-first species tree.** The tree was inferred using GeneRax and the RecPhyloxml output file was annotated. In this format, the gene tree is represented within the species tree. Major species groups are indicated on the right of the tree. Sub-lineages within the gene family are represented by coloured lines. Splits at species nodes represent speciation events. Gene duplications and losses are indicated with a square and a cross, respectively. A multi-pointed green star represents the duplication of origin of GCAP, that includes the ciliary phototransduction component. Other important duplication events are annotated throughout the tree.

**Supplementary Figure S23. Annotated GNB5 protein family tree reconciled with a ctenophore-first species tree.** The tree was inferred using GeneRax and the RecPhyloxml output file was annotated. In this format, the gene tree is represented within the species tree. Major species groups are indicated on the right of the tree. Splits at species nodes represent speciation events. Gene duplications and losses are indicated with a square and a cross, respectively. A green star represents the duplication of origin of the GNB5 family. Other important duplication events are annotated throughout the tree.

**Supplementary Figure S24. Annotated NCKX protein family tree reconciled with a ctenophore-first species tree.** The tree was inferred using GeneRax and the RecPhyloxml output file was annotated. In this format, the gene tree is represented within the species tree. Major species groups are indicated on the right of the tree. Sub-lineages within the gene family are represented by coloured lines. Splits at species nodes represent speciation events. Gene duplications and losses are indicated with a square and a cross, respectively. A multi-pointed green star represents the duplication of origin of NCKX 1/2, while a half-green, half-dark green star represents the duplication of origin of NCKX 3/4/5. Other important duplication events are annotated throughout the tree.

**Supplementary Figure S25. Annotated PDE6 A/B/C protein family tree reconciled with a ctenophore-first species tree.** The tree was inferred using GeneRax and the RecPhyloxml output file was annotated. In this format, the gene tree is represented within the species tree. Major species groups are indicated on the right of the tree. Sub-lineages within the gene family are represented by coloured lines. Splits at species nodes represent speciation events. Gene duplications and losses are indicated with a square and a cross, respectively. A multi-pointed green star represents the duplication of origin of PDE6 A/B/C, that includes the ciliary phototransduction components. Other important duplication events are annotated throughout the tree.

**Supplementary Figure S26. Annotated PDE6 G/H protein family tree reconciled with a ctenophore-first species tree.** The tree was inferred using GeneRax and the RecPhyloxml output file was annotated. In this format, the gene tree is represented within the species tree. Major species groups are indicated on the right of the tree. Splits at species nodes represent speciation events. Gene duplications and losses are indicated with a square and a cross, respectively. A multi-pointed green star represents the origin of the PDE6 G/H family. Other important duplication events are annotated throughout the tree.

**Supplementary Figure S27. Annotated recoverin protein family tree reconciled with a ctenophore-first species tree.** The tree was inferred using GeneRax and the RecPhyloxml output file was annotated. In this format, the gene tree is represented within the species tree. Major species groups are indicated on the right of the tree. Sub-lineages within the gene family are represented by coloured lines. Splits at species nodes represent speciation events. Gene duplications and losses are indicated with a square and a cross, respectively. A multi-pointed green star represents the duplication of origin of recoverin, that includes the ciliary phototransduction component.

**Supplementary Figure S28. Annotated RGS9/11 protein family tree reconciled with a ctenophore-first species tree.** The tree was inferred using GeneRax and the RecPhyloxml output file was annotated. In this format, the gene tree is represented within the species tree. Major species groups are indicated on the right of the tree. Sub-lineages within the gene family are represented by coloured lines. Splits at species nodes represent speciation events. Gene duplications and losses are indicated with a square and a cross, respectively. A multi-pointed green star represents the duplication of origin of RGS9/11, that includes the ciliary phototransduction component. Other important duplication events are annotated throughout the tree.

**Supplementary Figure S29. Annotated RGS9BP protein family tree reconciled with a ctenophore-first species tree.** The tree was inferred using GeneRax and the RecPhyloxml output file was annotated. In this format, the gene tree is represented within the species tree. Major species groups are indicated on the right of the tree. Sub-lineages within the gene family are represented by coloured lines. Splits at species nodes represent speciation events. Gene duplications and losses are indicated with a square and a cross, respectively. A half-green, half-dark green star represents the duplication of origin of RGS9BP, that includes the ciliary phototransduction component, and its paralog RGS7BP. Other important duplication events are annotated throughout the tree.

Supplementary Figure S30. Datasets used for the single-cell analyses. 12 species were used for the single-cell analyses. Phylogenetic relationships, as well as developmental stage, tissue type and source are indicated for each species. Focus was given on non-bilaterian species. Species silhouettes are images with CC0 1.0 Universal Public Domain Dedication licences obtained from <https://www.phylopic.org/>. Abbreviations: *D. mel*: *Drosophila melanogaster*; *H. sap*: *Homo sapiens*; *M. mus*: *Mus musculus*; *C. int*: *Ciona intestinalis*; *S. pur*: *Strongylocentrotus purpuratus*; *N. vec*: *Nematostella vectensis*; *S. pis*: *Stylophora pistillata*; *C. hem*: *Clytia hemisphaerica*; *H. vul*: *Hydra vulgaris*; *T. adh*: *Trichoplax adhaerens*; *A. que*: *Amphimedon queenslandica*; *M. lei*: *Mnemiopsis leidyi*.

**Supplementary Figure S31. Orthogroups of regulatory genes that are expressed in PRC-like metacells of at least two species.** The 286 orthogroups that are present in at least two species are listed in order of most frequent (with the hierarchy: present in most phyla, present in most species, present in most metacells). Species-specific orthogroups, present in two or more metacells, are not shown here but are listed in Supplementary Table S4. Expression in metacells is indicated with a black square when genes were amongst the top 100 differentially expressed genes of the metacell; while a grey square indicates expression with log-fold enrichment (lfp) > 0.5. Therefore, black squares indicate strong markers for a given metacell, while grey squares indicate that the gene is expressed in the metacell but differential expression level is not as high. Orthogroup names derive from the Preferred_names of the respective EggNog orthogroups, where present, or the EggNog orthogroup itself. GeneCards/Flybase were used to characterise the human/*Drosophila* representatives. Genes are highlighted: in bold if there is some evidence of involvement in vision and/or eye/photoreceptor development; in bold and underlined if there is strong evidence of involvement in vision and/or are expressed in the retina, although not necessarily in cones and rods; in bold, underlined and with asterisk if they are specifically expressed in photoreceptor cells. Species silhouettes were modified from images with CC0 1.0 Universal Public Domain Dedication licences obtained from <https://www.phylopic.org/>. Abbreviations: Dmel: *Drosophila melanogaster*; Hsap: *Homo sapiens*; Mmus: *Mus musculus*; Cint: *Ciona intestinalis*; Spur: *Strongylocentrotus purpuratus*; Nvec: *Nematostella vectensis*; Spis: *Stylophora pistillata*; Chem: *Clytia hemisphaerica*; Hvul: *Hydra vulgaris*; Tadh: *Trichoplax adhaerens*; Aque: *Amphimedon queenslandica*; Mlei: *Mnemiopsis leidyi*.

Supplementary Table Legends

Supplementary Table S1. All phototransduction components with respective gene and protein names. Common components are listed for both *Drosophila melanogaster* and *Homo sapiens*. Rhabdomeric components are listed for *D. melanogaster* and ciliary components are listed for *Homo sapiens*. The gene and protein names are based on FlyBase, GeneCards and UniProt.

Supplementary Table S2. List of eukaryotic species used for the phylogenetic analysis of phototransduction gene families. The source of the proteomes as well as the respective percentages of total complete BUSCO genes are indicated for each species. BUSCO was conducted using species proteomes versus the eukaryota_odb10 database.

**Supplementary Table S3. Comparison of the duplication and loss events in ctenophore-first versus sponge-first scenarios for each phototransduction gene family.** Sum of total events by number of sequences in the gene families is compared. Among the 29 gene families (the G alpha family is considered both as a whole and separated into rhabdomeric and ciliary subfamilies). 9 families show an identical number of events in both scenarios. Of the remaining 20 families, the ctenophore-first scenario has fewer events in a higher number of cases. Despite this, the absolute differences in total events between the two scenarios are minimal for all gene families with differences. This suggests that gene tree to species tree reconciliations yield similar results in terms of event counts for both alternative species tree scenarios.

**Supplementary Table S4. List of all EggNog orthogroups of regulatory genes and results of BLAST vs ATFDB.** Excel file containing: in sheet 1 the list of all 806 EggNog orthogroups of regulatory genes found throughout animal PRC-like metacells; in sheet 2 the 421 orthogroups present in at least two metacells with respective annotations and presence/absence in metacells; in sheet 3 results of BLAST of each orthogroup vs ATFDB; in sheet 4 some stats for the BLAST results.

**Supplementary Table S5. List of phototransduction gene families and respective keywords and profiles used for filtering.** Excel file containing: in sheet 1 the list of all phototransduction gene families with respective information about phototransduction pathway employed in as well as role in the cascade; in sheet 2 list of keywords used to filter sequences based on BLAST versus SwissProt database (if top 5 hits contained keywords then sequences were retained); in sheet 3 list of profiles used to retain sequences after InterProScan; in sheet 4 keywords and profiles for the whole G alpha family (as previous sheets separated G alpha q and G alpha i/o).

**Supplementary Table S6. List of evolutionary models used for each phototransduction gene family.** The model finder command used in IQTREE2 (with list of models tested) is indicated. For each gene family the result of Best-fit model based on BIC used for IQTREE2 trees and the best approximate model used for GeneRax are shown.

**Supplementary Table S7. Ranking system used to identify PRC-like metacells throughout animals.** For each of the 12 species for which single-cell data was analysed, expression of phototransduction genes was used to identify best candidate PRC-like metacells. In this excel file, for each species there is one sheet with the ranking calculations and one sheet with phototransduction genes differential expression levels (as identified by the log fold enrichment, lfp, defined in MetaCell) per metacells. Ranking was calculated by summing the scores from 7 categories: highest lfp for opsin(s); highest mean lfp of all phototranduction genes; highest mean lfp for common genes; highest mean lfp for ciliary or rhabdomeric genes (whichever is highest); highest percentage of total phototransduction genes; highest percentage of common phototransduction genes; highest percentage of either ciliary or rhabdomeric genes (whichever is highest).

**Supplementary Table S8. List of Pfam domains of known transcription factors used to identify regulatory genes orthogroups.** Candidate regulatory genes were scanned with InterProScan and when they had as hit one of the Pfam domains contained in this list, they were retained as candidate regulatory genes. This Method was complementary to using EggNog and keeping genes that fell in the COG category K, meaning that they are involved in transcription.
