## Supplementary figures and images for "A Pre-Bilaterian Origin of Phototransduction Genes and Photoreceptor Cells"

### FigureS1

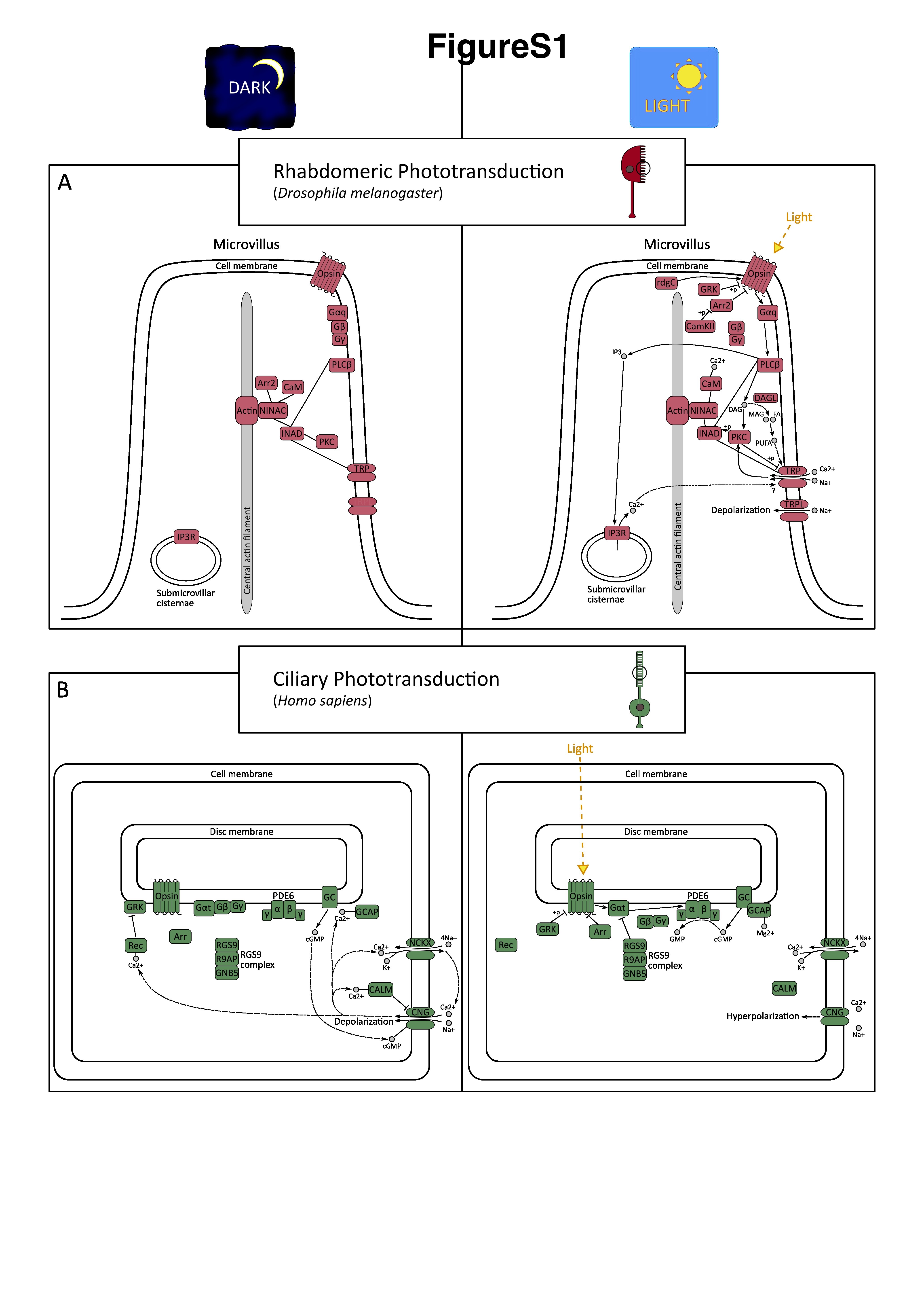

### FigureS2

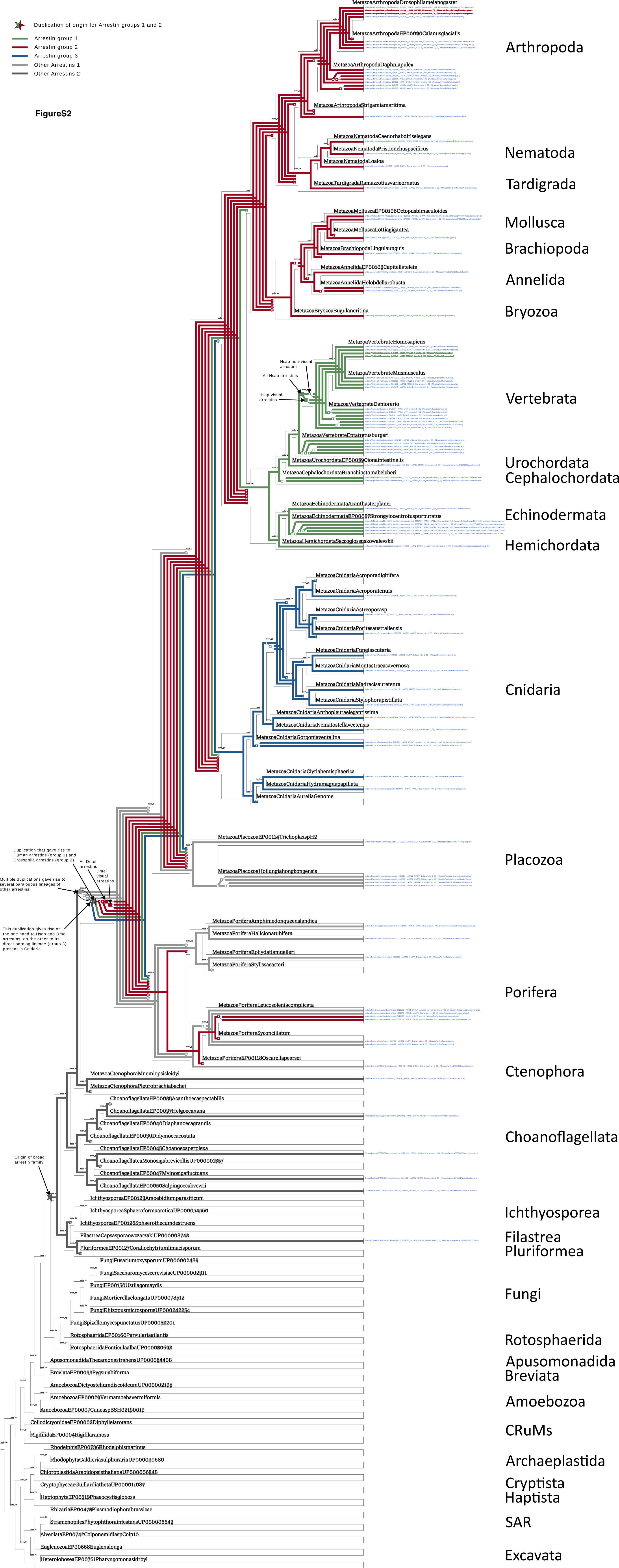

### FigureS5

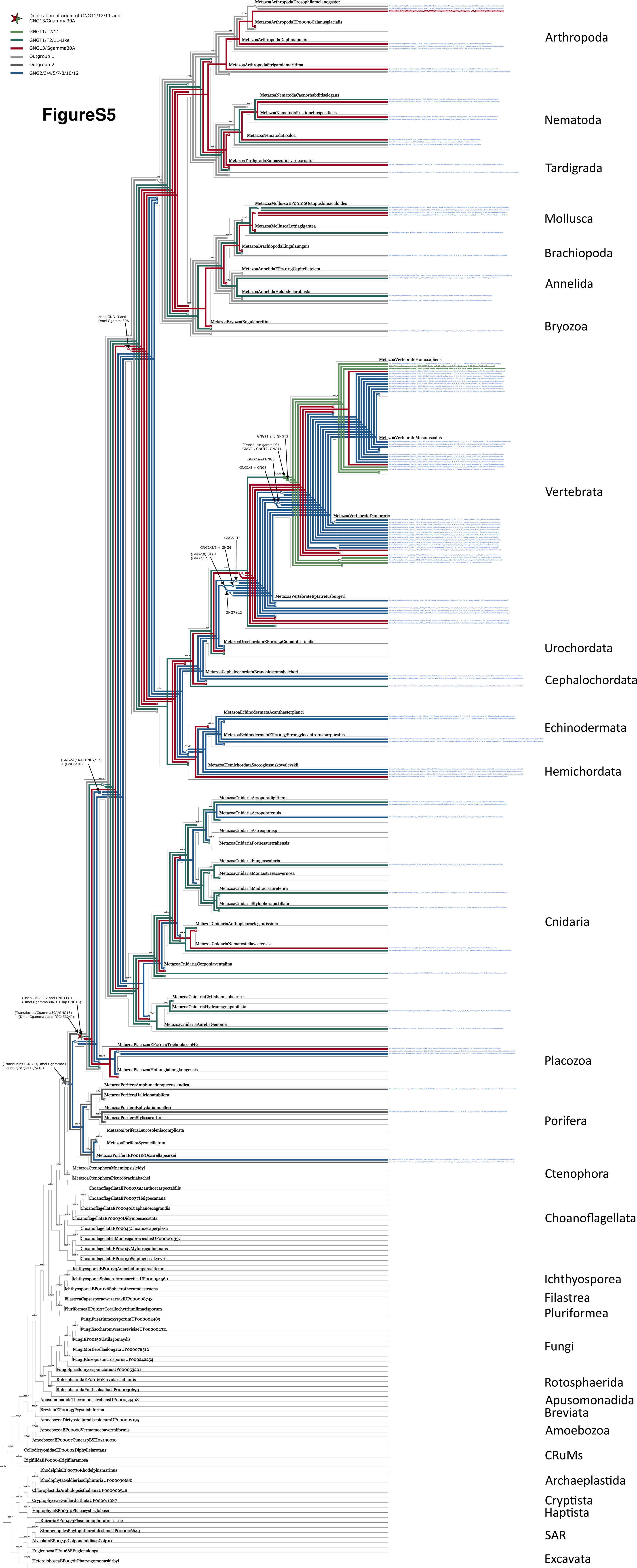

### FigureS6

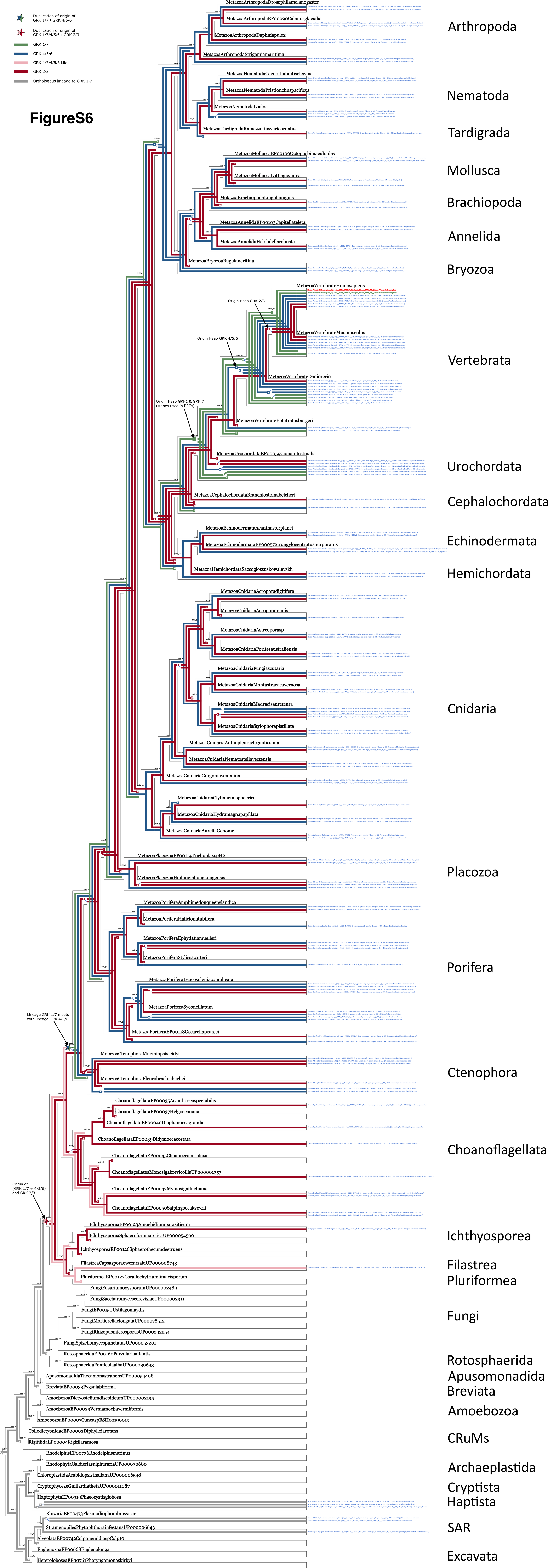

### FigureS9

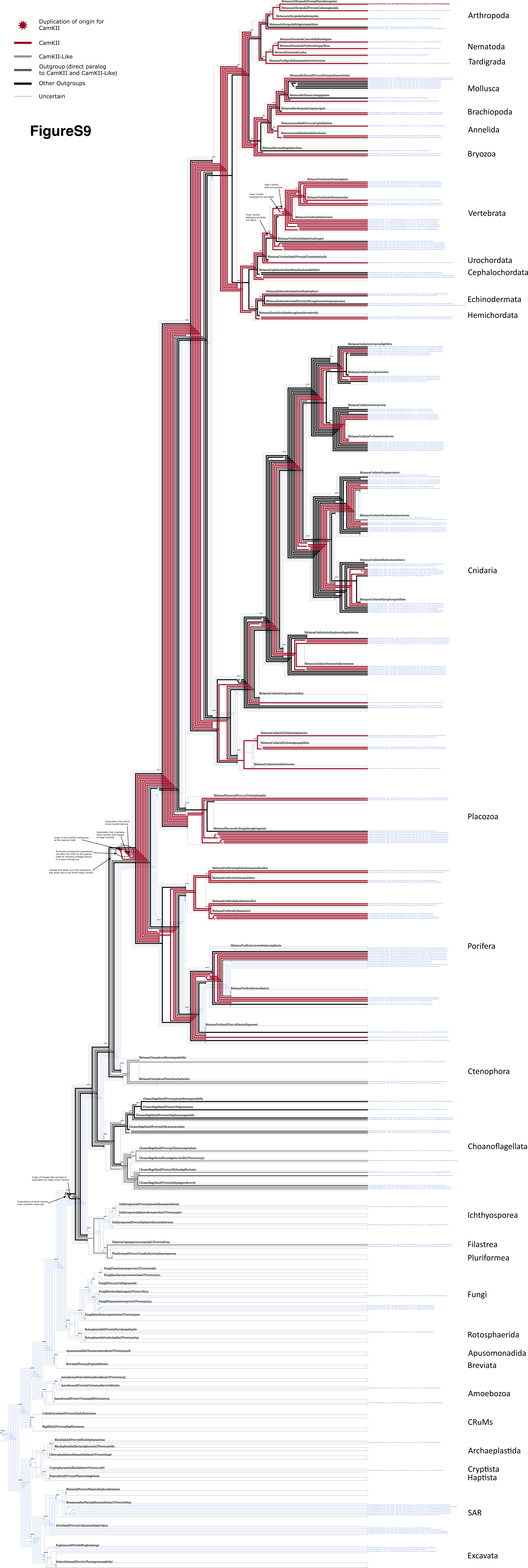

### FigureS10

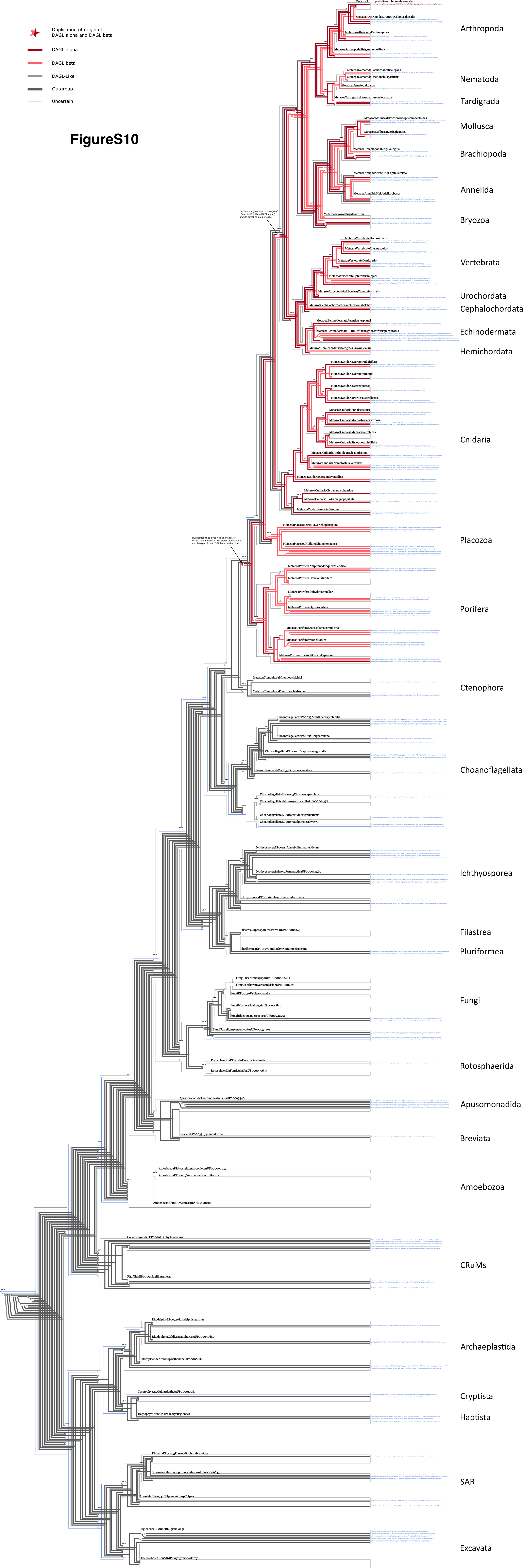

### FigureS11

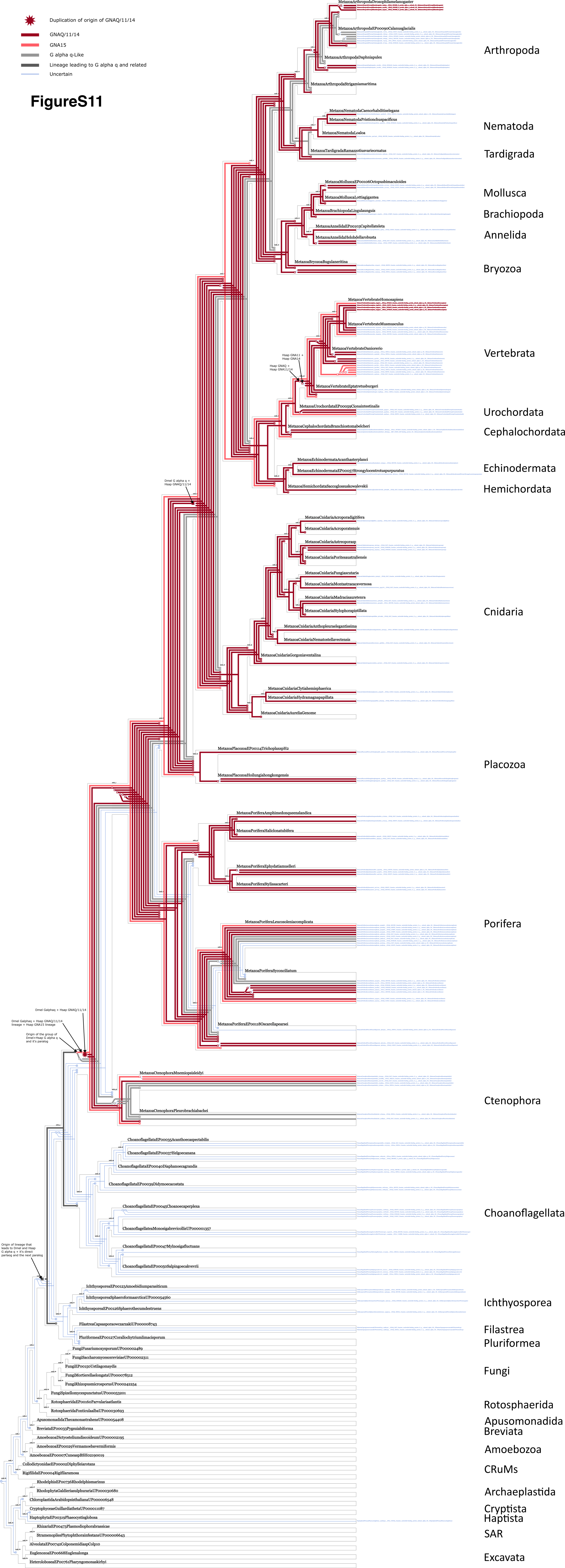

### FigureS12

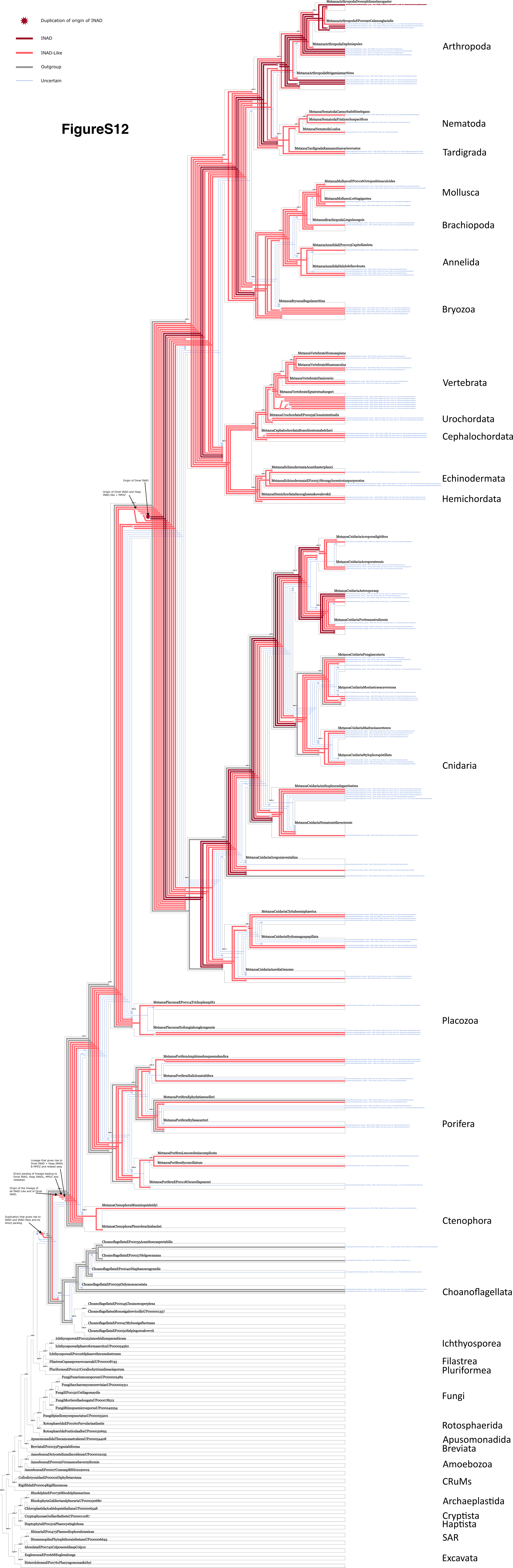

### FigureS13

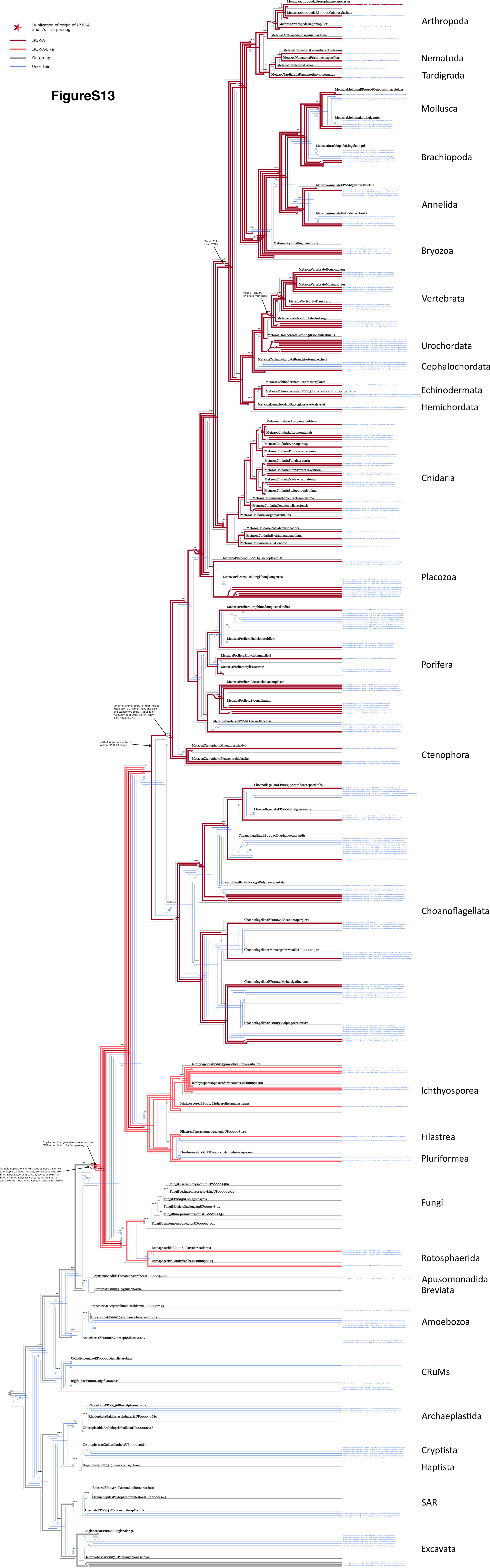

### FigureS17

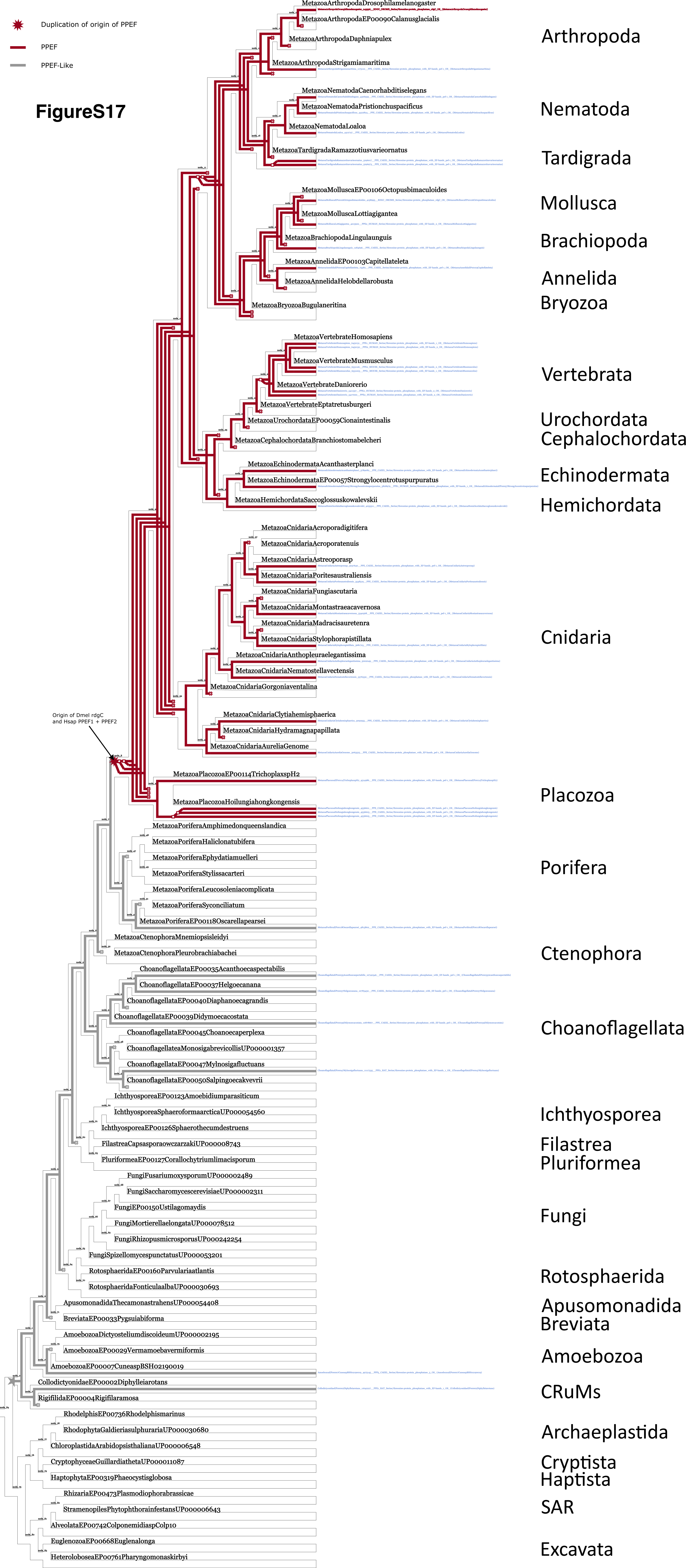

### FigureS18

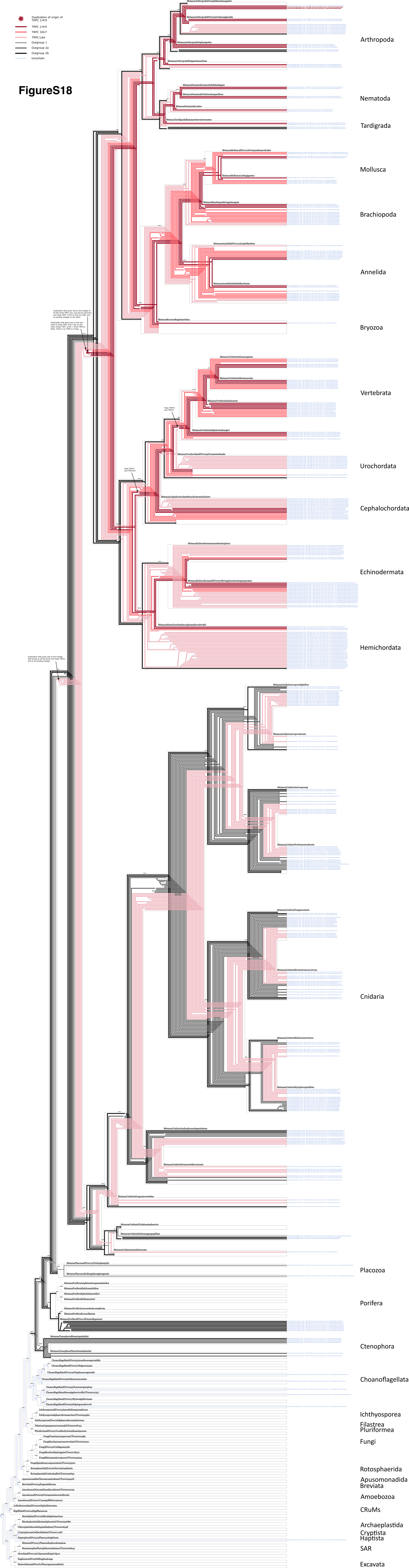

### FigureS20

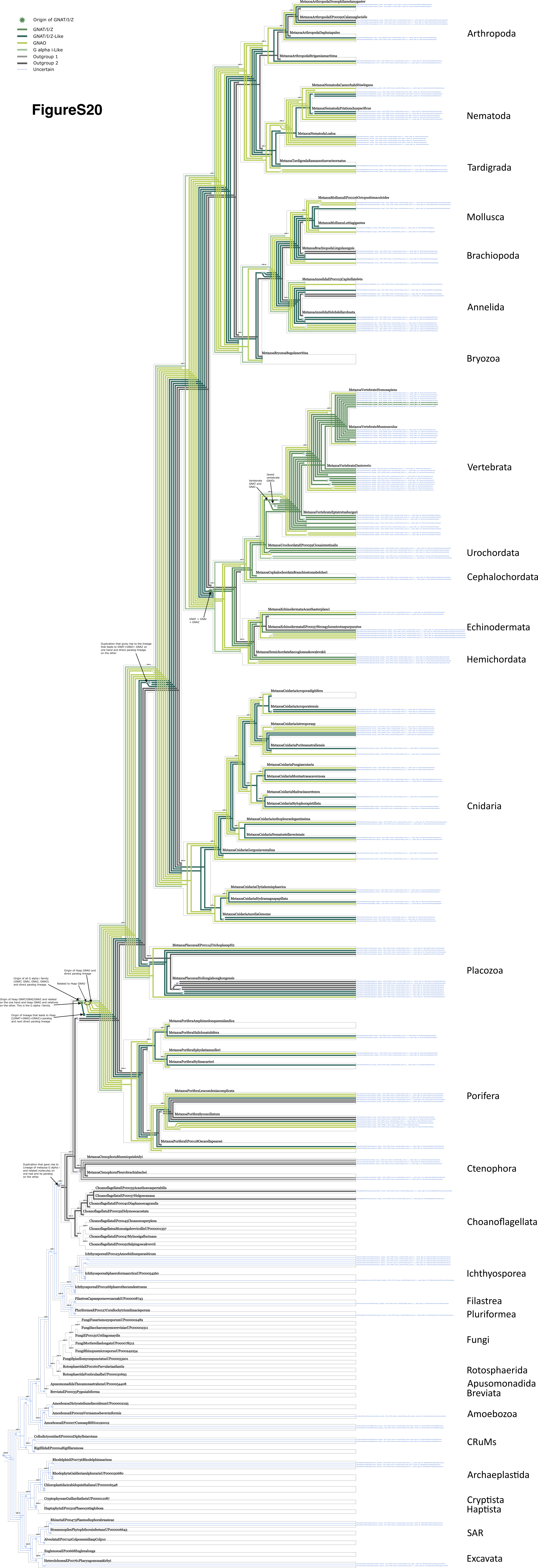

### FigureS21

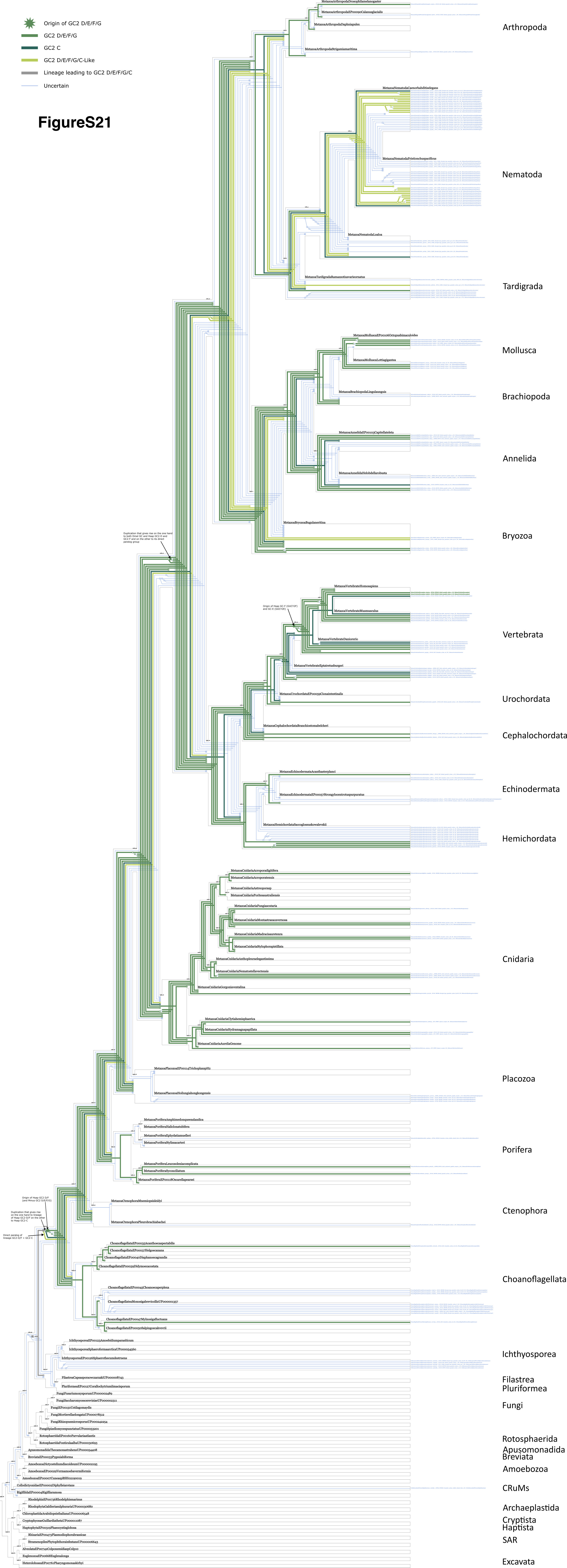

### FigureS22

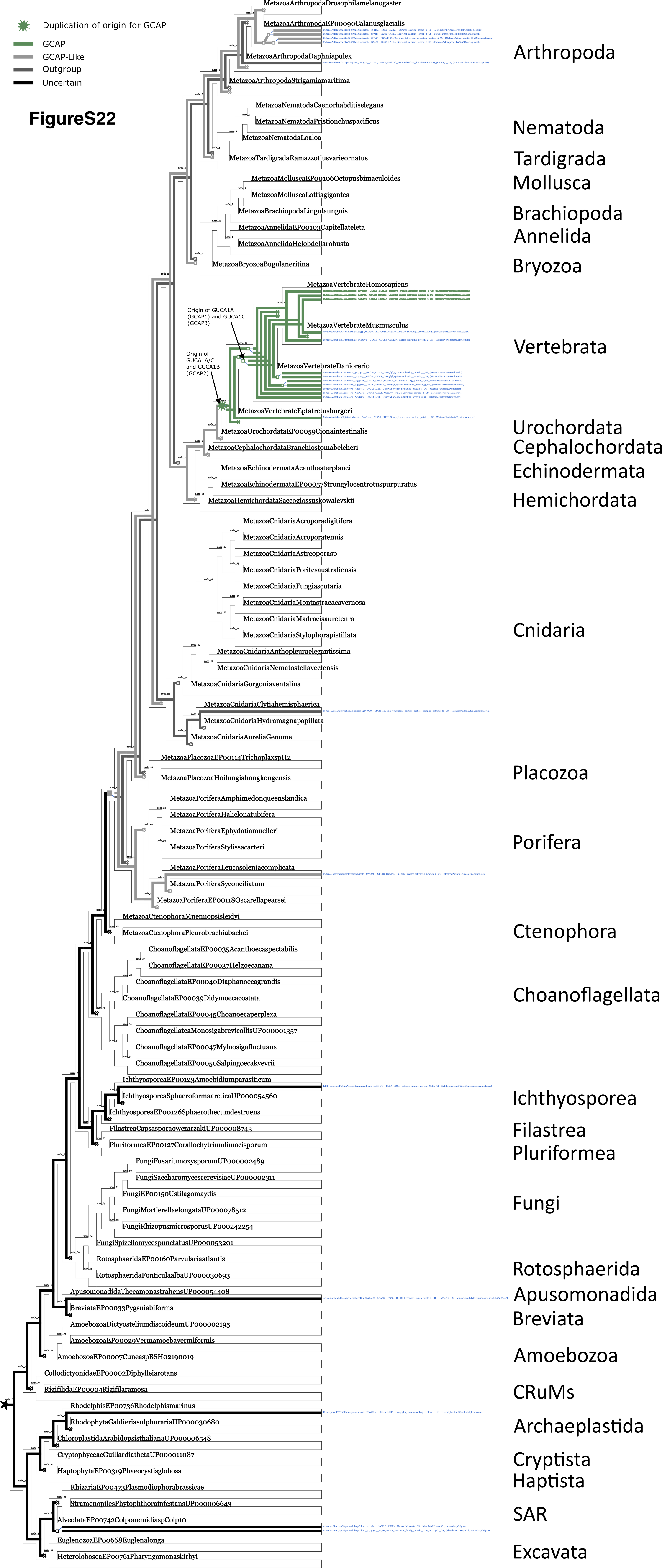

### FigureS23

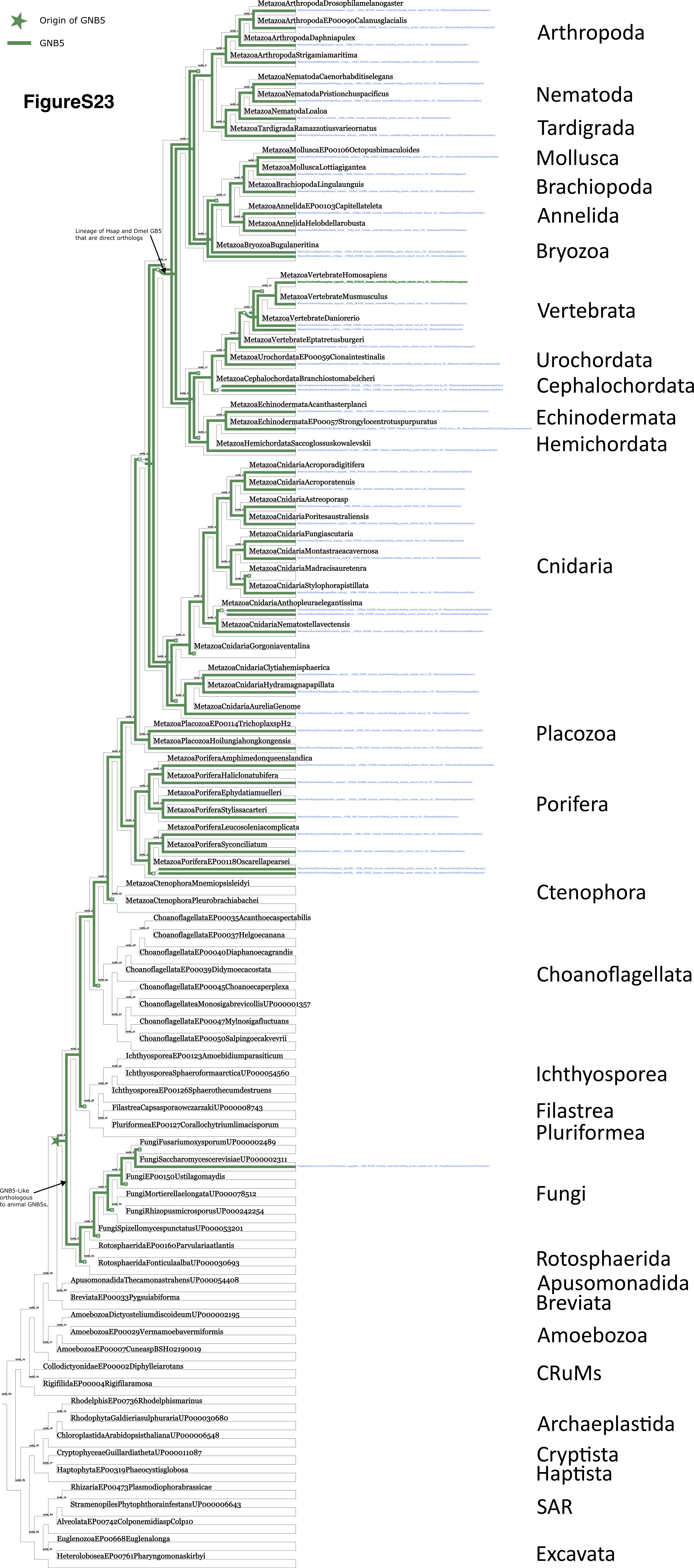

### FigureS25

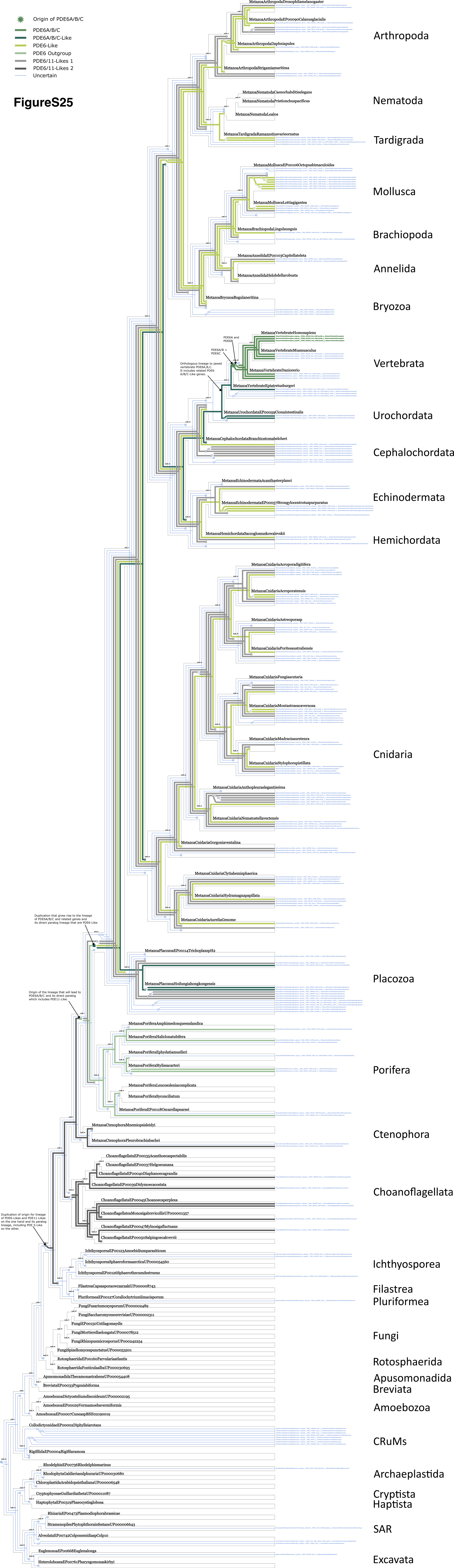

### FigureS26

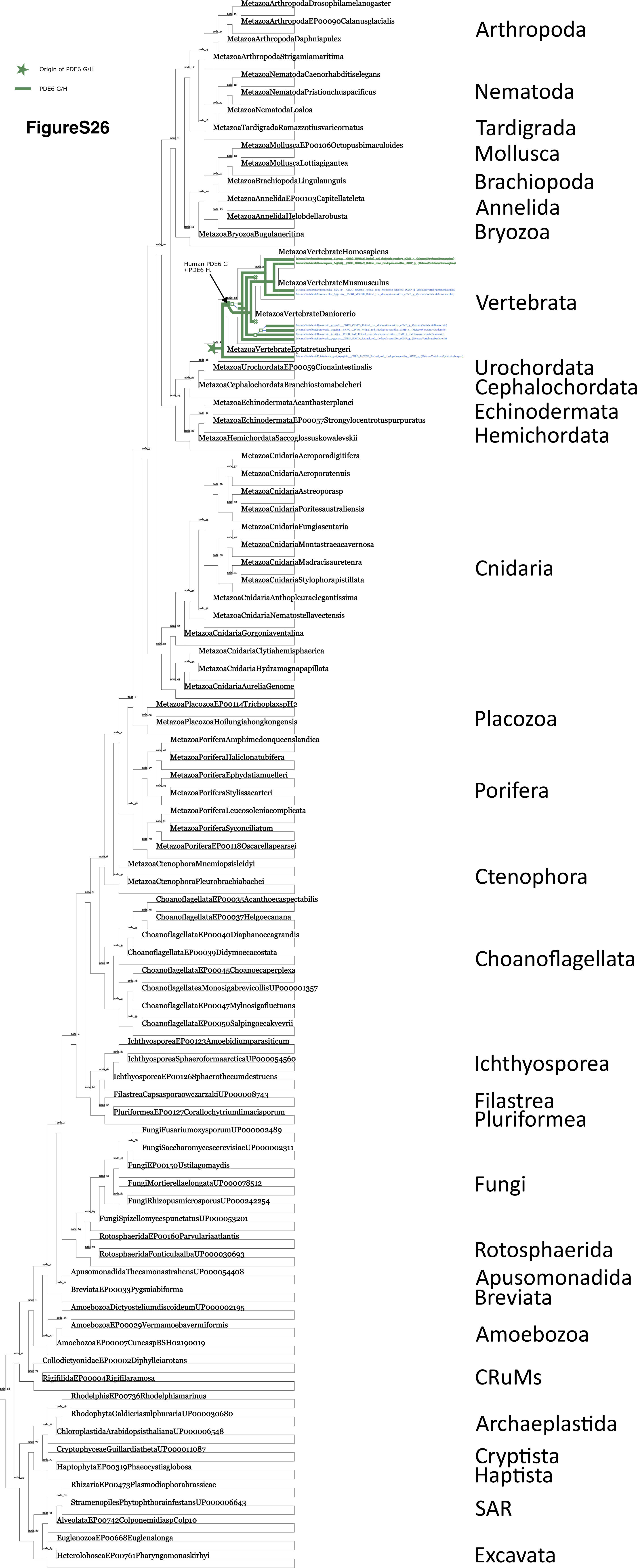

### FigureS27

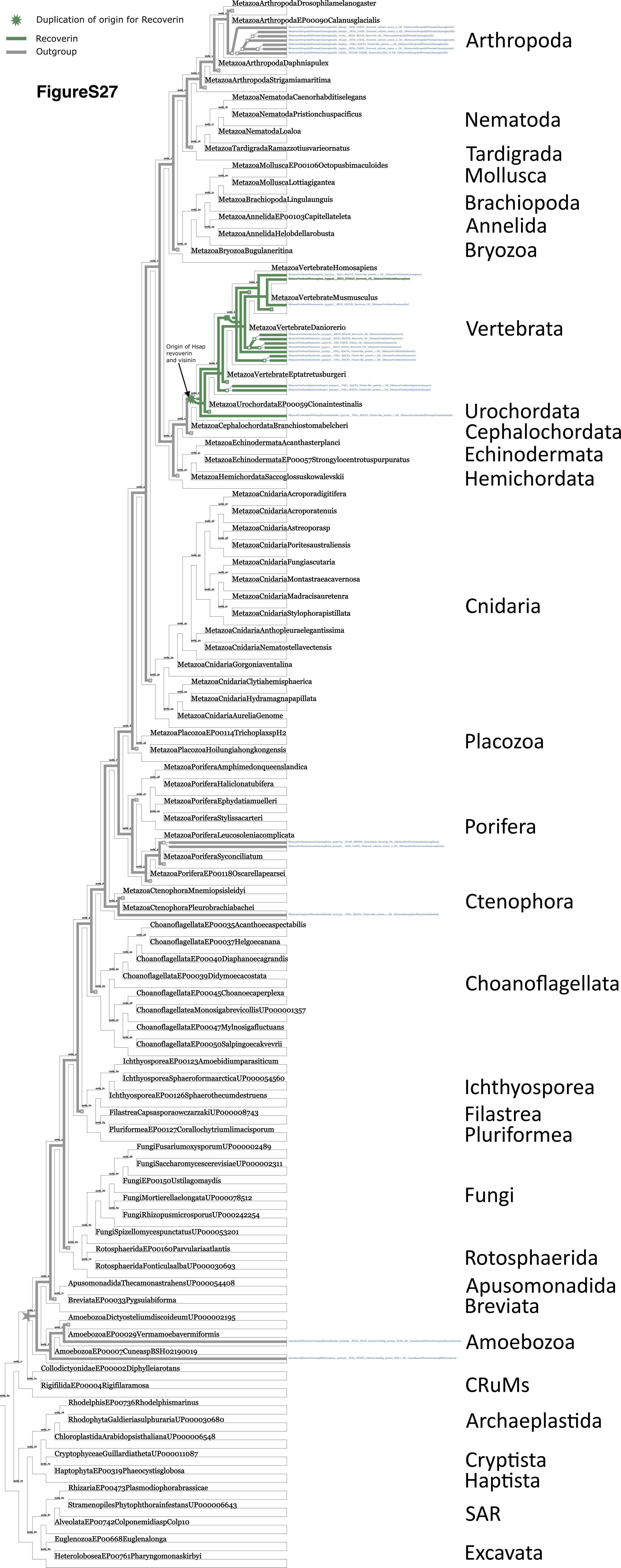

### FigureS28

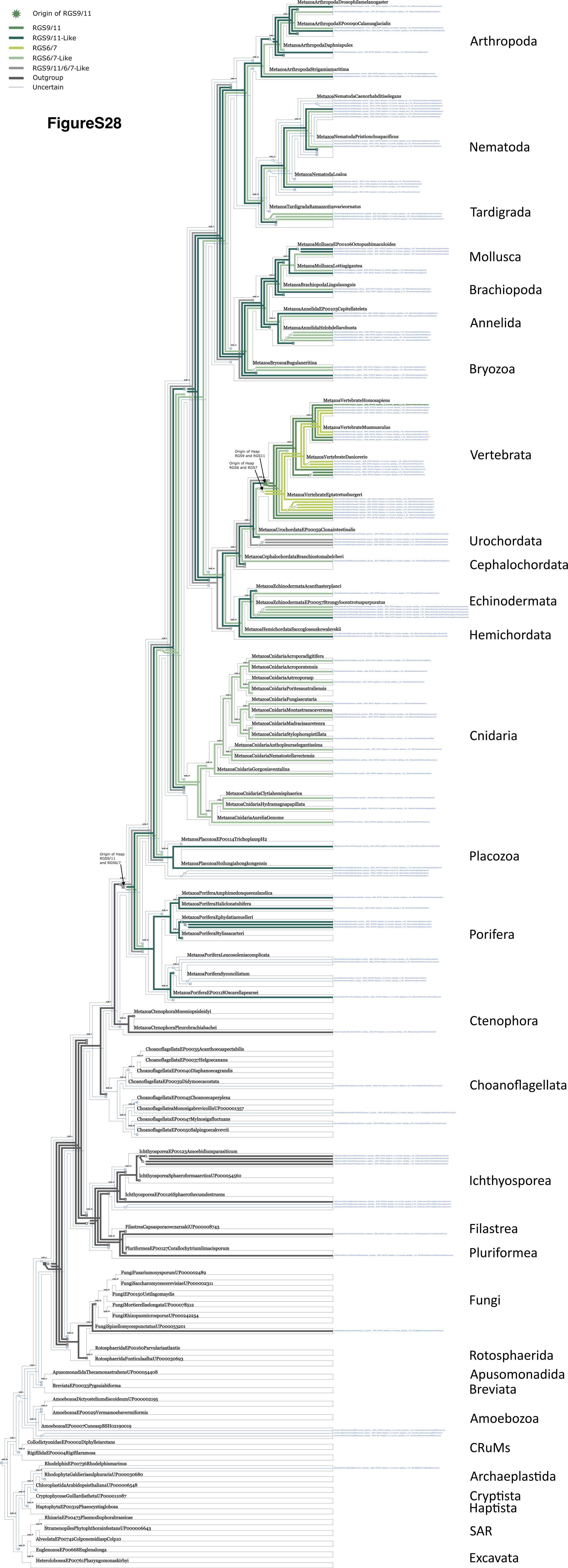

### FigureS29

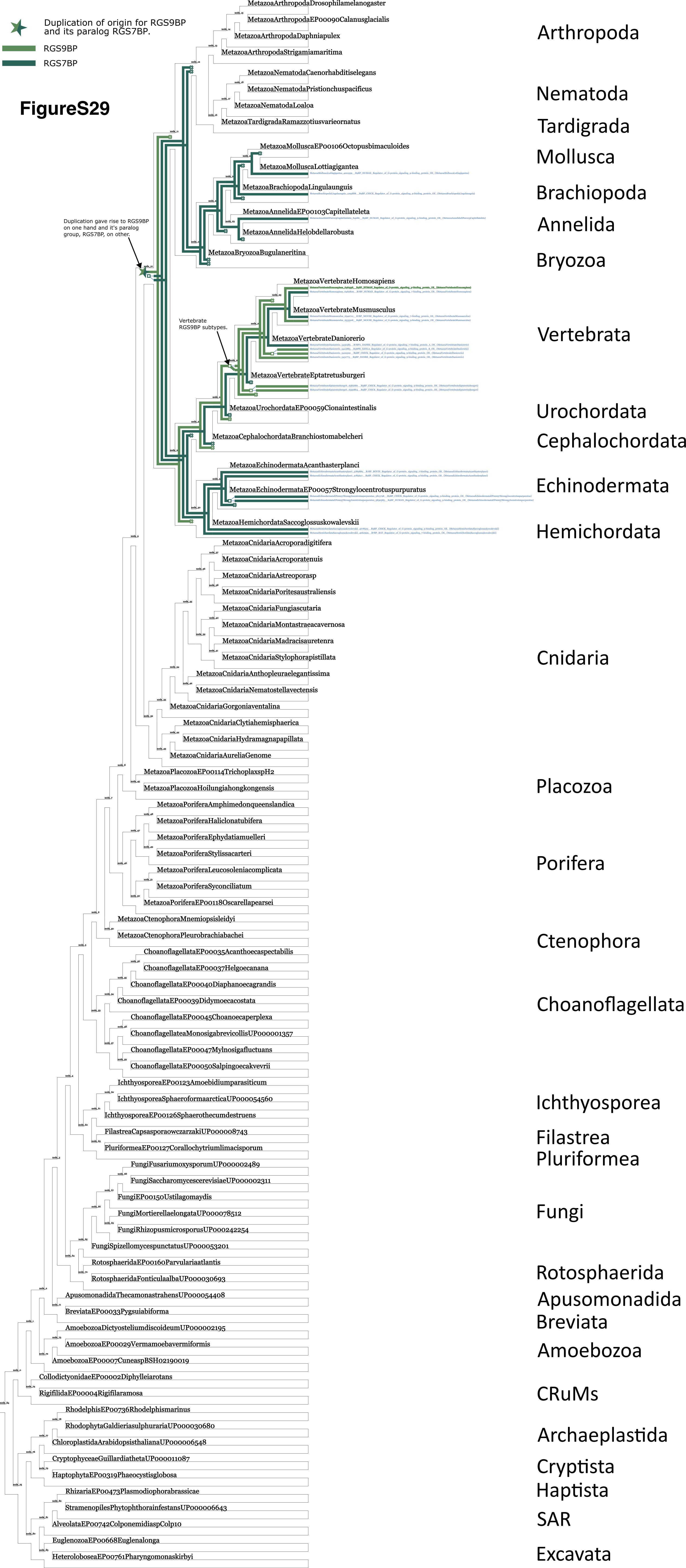

### FigureS30

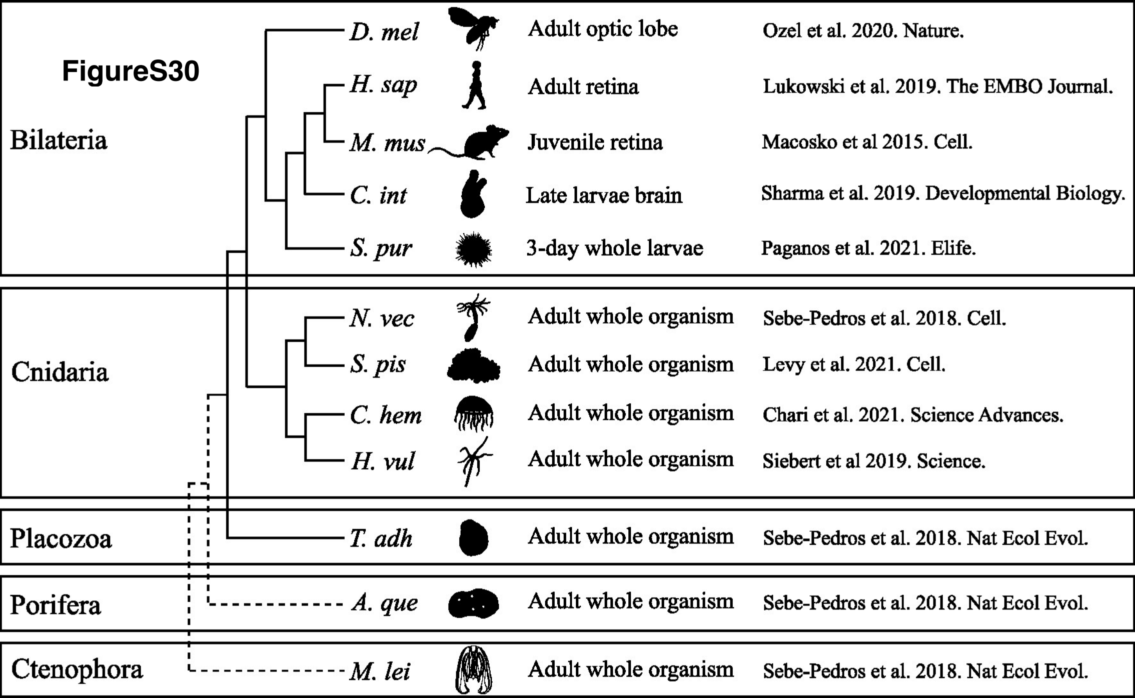

### FigureS31

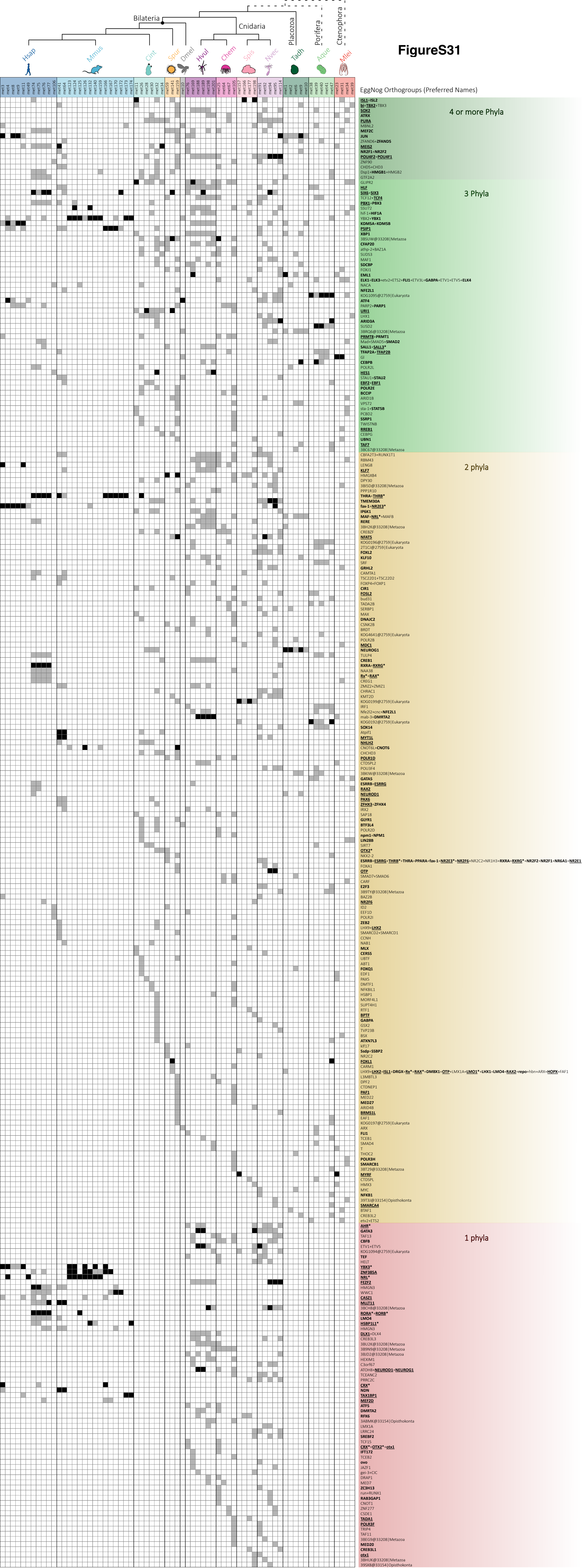
